# A community genomics approach to natural hybridization

**DOI:** 10.1101/2022.10.26.513841

**Authors:** Zachery D. Zbinden, Marlis R. Douglas, Tyler K. Chafin, Michael E. Douglas

## Abstract

Hybridization is a complicated, oft-misunderstood process. Once deemed unnatural and uncommon, hybridization is now recognized as ubiquitous among species. But hybridization rates within and among communities are poorly understood despite the relevance to ecology, evolution, and conservation. To clarify, we examined hybridization across 75 freshwater fish communities within the Ozarks of the North American Interior Highlands (USA) by SNP genotyping 33 species (*N*=2,865 individuals; ddRAD). We found evidence of hybridization (70 putative hybrids; 2.4% of individuals) among 18 species-pairs involving 73% (24/33) of study species, with the majority being concentrated within one family (Leuciscidae/minnows; 15 species; 66 individuals). Interspecific genetic exchange— or introgression— was evident from 24 backcrossed individuals (10/18 species-pairs). Hybrids occurred within 42 of 75 communities (56%). Four selected environmental variables (species richness, protected area extent, precipitation [May and annually]) exhibited 73–78% accuracy in predicting hybrid occurrence via random forest classification. Our community-level assessment identified hybridization as spatially widespread and environmentally dependent (albeit predominantly within one large, diverse family). Our approach provides a more holistic survey of natural hybridization by testing a wide range of species-pairs, thus contrasting with more conventional evaluations.

## 1. Introduction

Hybridization was once considered “exceedingly rare” [1], but it is now acknowledged as relatively common, primarily because of better detection using modern DNA sequencing [2,3]. A concurrent recognition is the significant role of hybridization and introgression in ecology and evolution [4–6], which diverges sharply from the historical perspective rooted in the biological species concept [7]. But while a growing catalog of species known to hybridize has helped to draw attention to hybridization (per-species documentation), the prevalence of hybrids within and among communities is poorly understood (per-individual documentation) [2].

The more frequently we encounter hybridization, the more evidence for its influential role in nature [2,8]. Hybridization and introgression are often considered maladaptive threats to biodiversity [6,9]. However, it is becoming more appreciated that the genetic novelty injected into a lineage through introgressive hybridization can be adaptive [8] and even provide evolutionary rescue [10]. Moreover, hybridization is expected to increase in lockstep with global environmental change [11,12]. Therefore, baseline estimates are needed to precisely gauge increases in hybridization and detect where it impacts ecosystems [10,13]. Attempts to understand the geographic patterns and processes of hybridization have not often considered an array of taxa. However, doing so will help identify where it occurs more generally and what environmental factors might drive it. These insights are essential because these places/environments may disproportionately impact biodiversity and ecosystems [14].

Broad comparative examinations often employ meta-analyses of numerous single-species-pair studies to quantify hybridization rates [15] or assess biogeographic relationships [14,16,17]. Several generalizations have emerged. Hybridization tends to be unevenly distributed across taxonomic groups and occurs more in plants than animals [2]. It has been documented more so in fish than other vertebrates [18], likely due to external fertilization within the aquatic environment [19]. Among fish, evidence of hybridization is more prevalent in freshwaters [1], especially between carps and minnows (Cypriniformes) [19], owing to their high diversity, sympatry, and breeding behaviors [20].

The less time since two species have diverged, the less time for pre- and post-zygotic barriers to evolve [20–22]. Similarly, the geographic overlap between two closely-related species also controls the pressure to evolve reproductive barriers [23], with most hybridizing lineages first evolving in allopatry [24]. Pre-zygotic barriers are often keyed (directly or indirectly) with the environment [25,26], weakening concurrently with environmental fluctuations, thus facilitating hybridization. For example, many fish rely on coloration to identify conspecific mates, and any environmental alterations impacting visual acuity (e.g., turbidity, siltation) can promote hybridization [27]. Much work has focused on human impacts due to environmental homogenization, sensory cue changes, and creating novel environments where hybrids may have advantages [28].

We quantified hybridization across a North American freshwater fish metacommunity by genotyping genome-wide single nucleotide polymorphisms (SNPs) that allowed us to detect hybrids among the individuals we sampled. We addressed the following questions: (i) How many hybrids occur within and among localities, and what is their frequency within species/families? (ii) Is genetic exchange among species (i.e., introgression) occurring as evinced by the presence of backcrossed individuals (e.g., *F*_1_ x Parental)? (iii) Is the incidence of hybrid individuals within communities predicated on environmental factors which allow the prediction of hybrid occurrence?

## 2. Methods

### (a) Data generation and processing

Our sampling area spanned the White River Basin (71,911 km^2^; Ozark Plateau, North American Interior Highlands). It represents an ideal study location because: (i) It is an unglaciated refugium with elevated fish diversity and limited anthropogenic impacts [29–31]; and (ii) Central to and representative of the Mississippi system, it is an excellent region from which generalizable patterns can be interpreted [32]. Sampling procedures were approved by the University of Arkansas Institutional Animal Care and Use Committee (IACUC #17077) and appropriate permitting agencies (Supplement S1). Fishes were seined (June 2017 to September 2018) and euthanized via tricaine methanesulfonate (MS-222) and 95% ethanol. Species diagnoses occurred in the laboratory.

Genomic DNA was isolated from fin clips (Qiagen Fast kits; Qiagen Inc.) and quantified by fluorometry (Qubit; Thermo-Fisher Scientific). Individuals were SNP genotyped using double-digest restriction site-associated DNA sequencing (ddRAD), using modified procedures [33,34], and sequenced as pooled batches (*N*=144/1×100 lane) on the Illumina HiSeq 4000 (Supplement S1).

Raw Illumina reads were demultiplexed (ipyrad; [35]), and family-level phylogenies were generated to verify individual species IDs via an alignment and assembly-free method (phyloRAD; [36]; Supplement S1). Individuals (*N*=3,042) were grouped by family (*N*=6) and processed de novo in ipyrad to generate separate family-level assemblies. Adapters/primers were removed, and reads with >5 low-quality bases (Phred<20) were discarded. Clusters were assembled using an 85% identity threshold, with loci subsequently removed via conditional criteria to ensure high-quality data (Supplement S1). Biallelic SNPs were further filtered and visualized using radiator [37] (Supplement S1).

### (b) Hybrid detection

Our initial objective was to detect putative hybrids and determine hybridization rates. We focused only within families for two reasons: No evidence implicates hybridization among these divergent families within six different orders; Also, a high number of species in RADseq alignments can reduce the power to detect hybrids by diminishing recovery of homologous loci (via allelic dropout from accumulating mutations in restriction enzyme cut-sites) [38]. All analyses were conducted using R 4.1.3 [39]. Individual genetic variation was first visualized using separate principal components analyses (PCAs) on each family-level SNP alignment (ade4; [40]).

Admixture analysis employing sparse non-negative matrix factorization (sNMF) was used to estimate ancestry coefficients for individuals within families (25 repetitions/*K* value (1–25) with regularization parameter (α)=100 (LEA; [41]). The best *K* from each run (via cross-validation) was used to impute missing data (*impute* function, method=‘mode’ in LEA), with sNMF repeated using imputed data (as above). An individual was flagged as a putative hybrid if the assignment probability for the majority ancestry cluster was <0.9.

Results were contrasted with a maximum-likelihood clustering approach considering hybrid categories per expected allele frequencies (Snapclust; [42]). Three models were tested: (i) *F*_1_ only; (ii) *F*_1_+ first-generation backcross; and (iii) *F*_1_+first- and second-generation backcrosses, with AIC used to determine the best fit [42]. Pairwise analyses were among species within families.

We assessed the power of sNMF and Snapclust to detect hybrids. We first removed putative hybrids following the above analyses. Then, we sampled alleles from the remaining pure individuals for each within-family species-pair to simulate hybridization between parentals (*P*_1_ & *P*_2_). We simulated 10 each of *F*_1_, *F*_2_, and backcross (5 *P*_1_ × *F*_1_ & 5 *P*_2_ × *F*_1_) (function *hybridize;* ADEGENET; [43]). SNPs for simulated hybrids were combined with parentals before running sNMF and Snapclust (as above). We then quantified the proportion of hybrids detected by each analysis.

### (c) Hybrid classification

Hybrids detected with sNMF and Snapclust were classified by assignment as either *F*_1_, *F*_2_, or backcross (NewHybrids; [44]). Backcrosses underscore hybrid viability and gene exchange among species (i.e., introgression). We implemented HybridDetective [45] to confirm sufficient statistical power for classification (*F*_1_, *F*_2_, or BXs). All NewHybrids analyses were conducted pairwise among species from which hybrids were detected. We first simulated known class hybrids by randomly sampling two alleles/locus from appropriate parental pools, with convergence assessed via three replicates. We then determined an optimal posterior probability threshold (=0.70) from which hybrid classes could be reliably assigned. The final MCMC was 1,000,000 iterations (250,000 burn-in). Assignments were made using a reduced panel of 200 SNPs exhibiting the greatest among-species differentiation (Weir and Cockerham’s *F*_ST_ among target species-pair) and lowest linkage disequilibrium (*r*^2^<0.2) via *getTopLoc*, HybridDetective [45].

### (d) Correlates of hybridization

We tested whether hybridizing/non-hybridizing species-pairs differed significantly with respect to genetic differentiation and basin-wide co-occurrence. Weir and Cockerham’s *F*_ST_ (diveRsity; [46]) and the number of co-occurrences across communities (cooccur; [47]) were calculated among *N*=137 species-pairs within families. Differences among hybridizing/non-hybridizing pairs were assessed using the Wilcoxon rank sum test.

We evaluated the influence of environmental factors on the incidence of hybrids within communities to determine their predictability. Each community (*N*=75) was classified as with/without hybrids, with potential predictor variables being: Species richness at each site; Drainage identity (8-digit USGS hydrologic unit); and high-resolution hydro-environmental descriptors (*N*=281, RiverATLAS; [48]). These encompassed factors broadly related to hydrology, physiography, climate, land cover, geology, and anthropogenic impact. Associations between hybrid incidence and predictive variables were assessed using random forest classification (i.e., supervised machine-learning employing linear/non-linear relationships among mixed data types [49]). Bootstrapped decision trees (via predictor variables) are aggregated, with each trained on a random 2/3 of samples (=“in-bag”), with validation via the remaining 1/3 (=“out-of-bag”).

We implemented the random forest quantile-classifier approach [50] in randomForestSRC [51] using 100,000 bootstrapped decision trees (=ntree), each of which randomly sampled 1/3 of total predictors (=mtry). Variable Importance (VIMP; Increase in prediction error when a variable is randomized) was quantified via permutation for each predictive variable. VIMPs significantly >0 were assessed by subsampling 1,000 trees. Random forest variable selection was executed using minimal depth and a conservative threshold set to “high,” employing only that subset of predictive variables with significant VIMP [52]. A final random forest model was built from this selected subset of predictors. Accuracy was evaluated using *G*-mean (0–1; the geometric mean of the true negative and positive rates), which replaces the misclassification rate in imbalanced data settings [50] and the normalized Brier score (0–1; mean square difference between true classes and predicted probabilities).

## 3. Results

Fish collections across 75 communities/sites yielded 72 species (*N*=3,605 individuals), averaging 10.8 species/site—typical for streams across the Mississippi Basin [53]. We genotyped 33 species to represent the fish metacommunity [54] (i.e., 84% of total individuals collected, averaging 9.44 spp./site). Of *N*=137 intra-familial species-pairs examined, 13 (9.5%) did not co-occur at our sites. Four do have non-overlapping, parapatric distributions, each with one occurring in the upper White and the other in the Black River: one ictalurid (*Noturus albater/ maydeni*); one leuciscid (*Luxilus pilsbryi/zonatus*); and two percids (*Etheostoma juliae/ uniporum; E. spectabile/ uniporum*). Despite range overlap, nine other pairs did not co-occur (six leuciscids and three percids), which may be due to chance/sampling, given that pairs included at least one uncommon species (≤10 occurrences).

We examined *N*=2,865 individuals across six families post-filtering using SNP genotypes (Table 1; mean missing data=21%; mean coverage=56x). SNPs varied by family and were inversely related to the number of species (Table 1). Power analyses verified the robustness of panels for detecting/classifying hybrids based on genotype frequencies (Supplement S2–S19). Among *N*=30 simulated hybrids for each unique species-pair assessed (*N*=137), sNMF and Snapclust detected 93% and 100%, respectively. Simulated hybrids among pairs targeted for classification via HybridDetective and NewHybrids showed 90-100% assignment accuracy across genotype frequency classes and species-pairs for posterior probability thresholds ≥ 0.70.

**TABLE 1.** Samples (N=33 species) from the White River Basin, USA. *N_I_*=number of individuals/species (post-filtering), *N_S_* =number of collection sites/species (post-filtering), Miss=mean proportion of missing data, Depth=mean sequencing depth, *H_o_*=observed heterozygosity, *N_FA_*=number of individuals/family, SNPs=single nucleotide polymorphisms in family panel (post-filtering).

| Family | Common Name | Scientific Name | $N_I$ | $N_S$ | Miss | Depth | $H_o$ | $N_{FA}$ | SNPs |
| --- | --- | --- | --- | --- | --- | --- | --- | --- | --- |
| Centrarchidae | Bluegill Sunfish | *Lepomis macrochirus* | 69 | 23 | 0.15 | 46 | 0.028 | 375 | 1926 |
|  | Longear Sunfish | *Lepomis megalotis* | 239 | 54 | 0.09 | 47 | 0.036 |  |  |
|  | Smallmouth Bass | *Micropterus dolomieu* | 43 | 30 | 0.47 | 39 | 0.011 |  |  |
|  | Largemouth Bass | *Micropterus salmoides* | 24 | 17 | 0.46 | 41 | 0.013 |  |  |
| Cottidae | Banded Sculpin | *Cottus caroliniae* † | 33 | 18 | 0.08 | 64 | 0.043 | 75 | 5344 |
|  | Ozark Sculpin | *Cottus hypselurus* ‡ | 42 | 10 | 0.16 | 63 | 0.057 |  |  |
| Fundulidae | Northern Studfish | *Fundulus catenatus* | 105 | 28 | 0.12 | 37 | 0.008 | 226 | 2366 |
|  | Blackspotted Topminnow | *Fundulus olivaceus* | 121 | 34 | 0.20 | 33 | 0.024 |  |  |
| Ictaluridae | Ozark Madtom | *Noturus albater* | 10 | 5 | 0.12 | 65 | 0.033 | 31 | 2744 |
|  | Slender Madtom | *Noturus exilis* | 16 | 12 | 0.09 | 51 | 0.042 |  |  |
|  | Black River Madtom | *Noturus maydeni* | 5 | 3 | 0.28 | 34 | 0.034 |  |  |
| Leuciscidae | Central Stoneroller | *Campostoma anomalum* | 128 | 44 | 0.27 | 58 | 0.010 | 1507 | 343 |
|  | Largescale Stoneroller | *Campostoma oligolepis* | 135 | 41 | 0.29 | 52 | 0.008 |  |  |
|  | Southern Redbelly Dace | *Chrosomus erythrogaster* | 38 | 10 | 0.41 | 38 | 0.001 |  |  |
|  | Whitetail Shiner | *Cyprinella galactura* | 75 | 16 | 0.19 | 59 | 0.012 |  |  |
|  | Steelcolor Shiner | *Cyprinella whipplei* | 30 | 8 | 0.27 | 65 | 0.010 |  |  |
|  | Striped Shiner | *Luxilus chrysocephalus* | 63 | 18 | 0.18 | 59 | 0.013 |  |  |
|  | Duskystrip Shiner | *Luxilus pilsbryi* | 258 | 33 | 0.09 | 76 | 0.023 |  |  |
|  | Bleeding Shiner | *Luxilus zonatus* | 100 | 17 | 0.09 | 85 | 0.013 |  |  |
|  | Redfin Shiner | *Lythrurus umbratilis* | 24 | 5 | 0.26 | 47 | 0.004 |  |  |
|  | Bigeye Shiner | *Notropis boops* | 226 | 31 | 0.10 | 89 | 0.019 |  |  |
|  | Ozark Minnow | *Notropis nubilus* | 193 | 35 | 0.11 | 76 | 0.016 |  |  |
|  | Carmine Shiner | *Notropis percobromus* | 67 | 15 | 0.20 | 66 | 0.010 |  |  |
|  | Telescope Shiner | *Notropis telescopus* | 83 | 15 | 0.14 | 75 | 0.006 |  |  |
|  | Bluntnose Minnow | *Pimephales notatus* | 55 | 24 | 0.25 | 73 | 0.007 |  |  |
|  | Creek Chub | *Semotilus atromaculatus* | 32 | 14 | 0.32 | 51 | 0.001 |  |  |
| Percidae | Greenside Darter | *Etheostoma blennioides* | 62 | 26 | 0.24 | 55 | 0.006 | 651 | 687 |
|  | Rainbow Darter | *Etheostoma caeruleum* | 348 | 53 | 0.09 | 54 | 0.026 |  |  |
|  | Fantail Darter | *Etheostoma flabellare* | 26 | 11 | 0.33 | 38 | 0.001 |  |  |
|  | Yoke Darter | *Etheostoma juliae* § | 63 | 15 | 0.24 | 58 | 0.011 |  |  |
|  | Orangethroat Darter | *Etheostoma spectabile* ¶ | 50 | 10 | 0.24 | 48 | 0.023 |  |  |
|  | Current Darter | *Etheostoma uniporum* | 18 | 7 | 0.25 | 41 | 0.005 |  |  |
|  | Banded Darter | *Etheostoma zonale* | 84 | 26 | 0.25 | 51 | 0.018 |  |  |

Note: Several taxonomic changes have been adopted in the most recent *Fishes of Arkansas* [30] based on the literature but have yet to be recognized by the American Fisheries Society or Eschmeyers's Catalog of Fishes.
† *Uranidea caroliniae*
‡ *Uranidea immaculata* (Knobfin Sculpin)
§ *Nothonotus juliae*
¶ *Etheostoma* sp. cf. *spectabile* (Ozark Darter)

### (a) Hybrid detection

The number of species-pairs examined varied considerably among the six families/orders, with just one pair tested for two families versus 105 pairs within Leuciscidae (Table 2). Hybrids (*N*=70; 2.4%) were detected within four families (Table 2), with hybrid proportions ranging from 0–4.4% of individuals. All but four hybrids were minnows (Leuciscidae), with exceptions being: (i) *Micropterus dolomieu* x *salmoides* (Centrarchidae); (ii) *Etheostoma juliae* x *zonale* (Percidae); (iii) *Etheostoma spectabile* x *caeruleum* (Percidae); and (iv) a putative multi-specific hybrid: *Noturus maydeni* x *albater* x *exilis* (Ictaluridae). Thus, we recognized *N*=18 hybridizing species-pairs and *N*=8 species-triplets, i.e., multi-species hybrids. We did not detect hybrids within Cottidae or Fundulidae.

**TABLE 2.** Hybridization within six families sampled across the White River Basin, USA. Columns include: N Species=number of species analyzed; Unique pairs=number of unique intra-familial; Pairs Not Co-occurring=number of pairs within a family not found together at the same site; N Indiv=number of individuals examined; N Hybrids=number of hybrids detected; Percent Indiv=Hybrid percentage. Unique species-pairs with hybrids are only considered for ancestry between two species (i.e., species triplets were not counted).

| Family | N Species | Unique Pairs | Pairs Not Co-Occurring | N Indiv. | N Hybrids | Percent Indiv. | Species w/ Hybrids | Percent Species w/ Hybrid | Unique Pairs w/ Hybrid | Percent Pairs w/ Hybrid |
| --- | --- | --- | --- | --- | --- | --- | --- | --- | --- | --- |
| Fundulidae | 2 | 1 | 0 | 226 | 0 | 0.0% | 0 | 0% | 0 | 0% |
| Cottidae | 2 | 1 | 0 | 75 | 0 | 0.0% | 0 | 0% | 0 | 0% |
| Ictaluridae | 3 | 3 | 1 | 31 | 1 | 3.2% | 3 | 100% | 0 | 0% |
| Centrarchidae | 4 | 6 | 0 | 375 | 1 | 0.3% | 2 | 50% | 1 | 17% |
| Percidae | 7 | 21 | 5 | 651 | 2 | 0.3% | 4 | 57% | 2 | 10% |
| Leuciscidae | 15 | 105 | 7 | 1507 | 66 | 4.4% | 15 | 100% | 15 | 14% |
| Overall | 33 | 137 | 13 | 2865 | 70 | 2.4% | 24 | 73% | 18 | 13% |

Within Leuciscidae, we identified 66 hybrids and 15 hybridizing species-pairs (Supplement S20). The most remarkable (*N*=29 individuals; 41%) involved *Campostoma anomalum* x *oligolepis*. All minnow species herein (*N*=15) appear to hybridize with at least one other species in the study basin. The number of hybrids per species was significantly related to the number of community occurrences (*R^2^*=0.16, *F*=6.1, *p*=0.02).

PCA, sNMF, and Snapclust were largely congruent in detecting hybrids (Figures 1–2). Both sNMF and Snapclust detected the same 36 hybrids, each separately identifying additional (26 and 8, respectively). Snapclust showed excellent power in detecting hybrids based on allele frequencies (100%). However, it comes with a cost: the same individual was often suggested as a hybrid in multiple pairwise tests, regardless of the second parental species. Fortunately, we could adjudicate these using PCA and sNMF. A list of hybrid individuals and inferences for each are available in Supplementary Material (S21).

**FIGURE 1.**
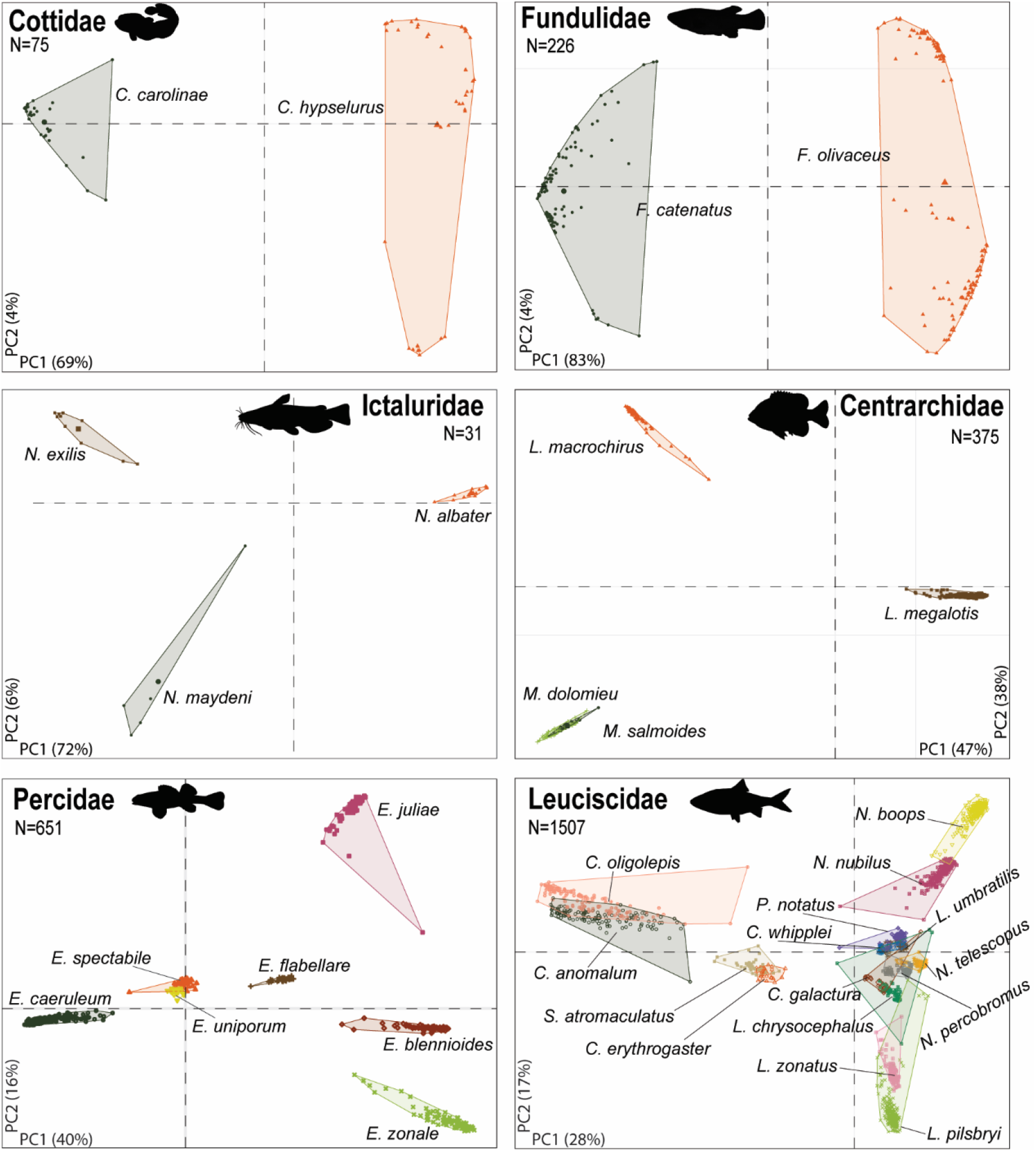
First two principal components (PCs) derived from SNP genotypes in six stream fish families collected across the White River Basin, USA. The variance explained by each component is in the bottom left or right corner of each plot. *N*=numbers of individuals/family.

**FIGURE 2.**
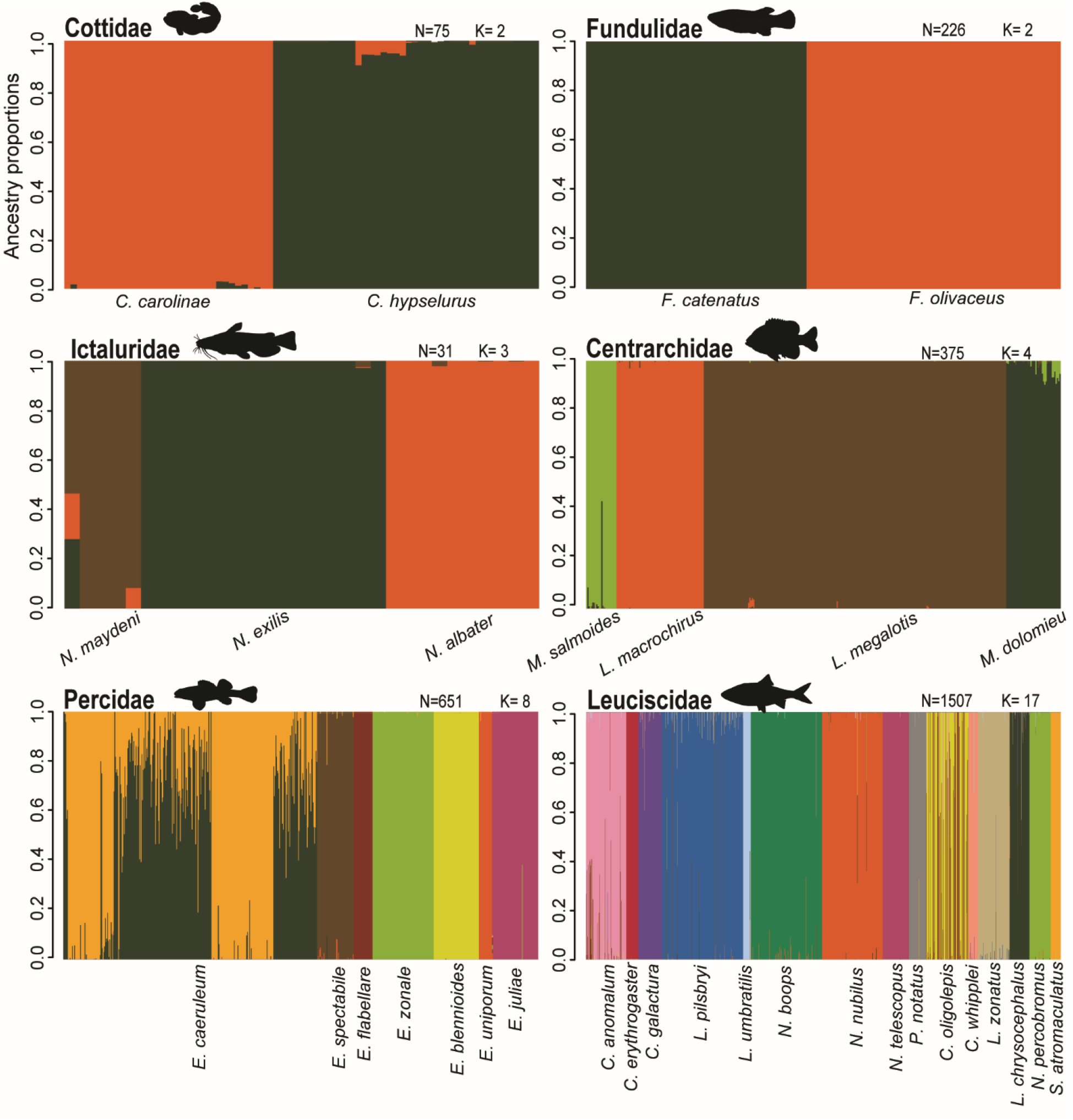
Calculated ancestral proportions for *N*=2,865 individuals from *N*=33 fish species grouped by family collected from the White River Basin, USA. *N*=number of individuals, *K*=optimum number of clusters.

### (b) Hybrid classification and introgression

We also found evidence of introgression based on the occurrence of backcrossed hybrid individuals. NewHybrids identified 39% of hybrids as backcrossed (24/62, excluding eight multi-specific hybrids); this corresponds to 55% (10/18) of all putatively hybridizing species-pairs showing evidence of introgressive hybridization (Table 3). Approximately 24% of hybrids (15/62) were seemingly *F*_1_ or *F*_2_. Finally, NewHybrids designated 37% (23/62) of putative hybrids as pure parentals, although possibly introgressed individuals poorly classified, i.e., late-generation hybrids [42].

**TABLE 3.**
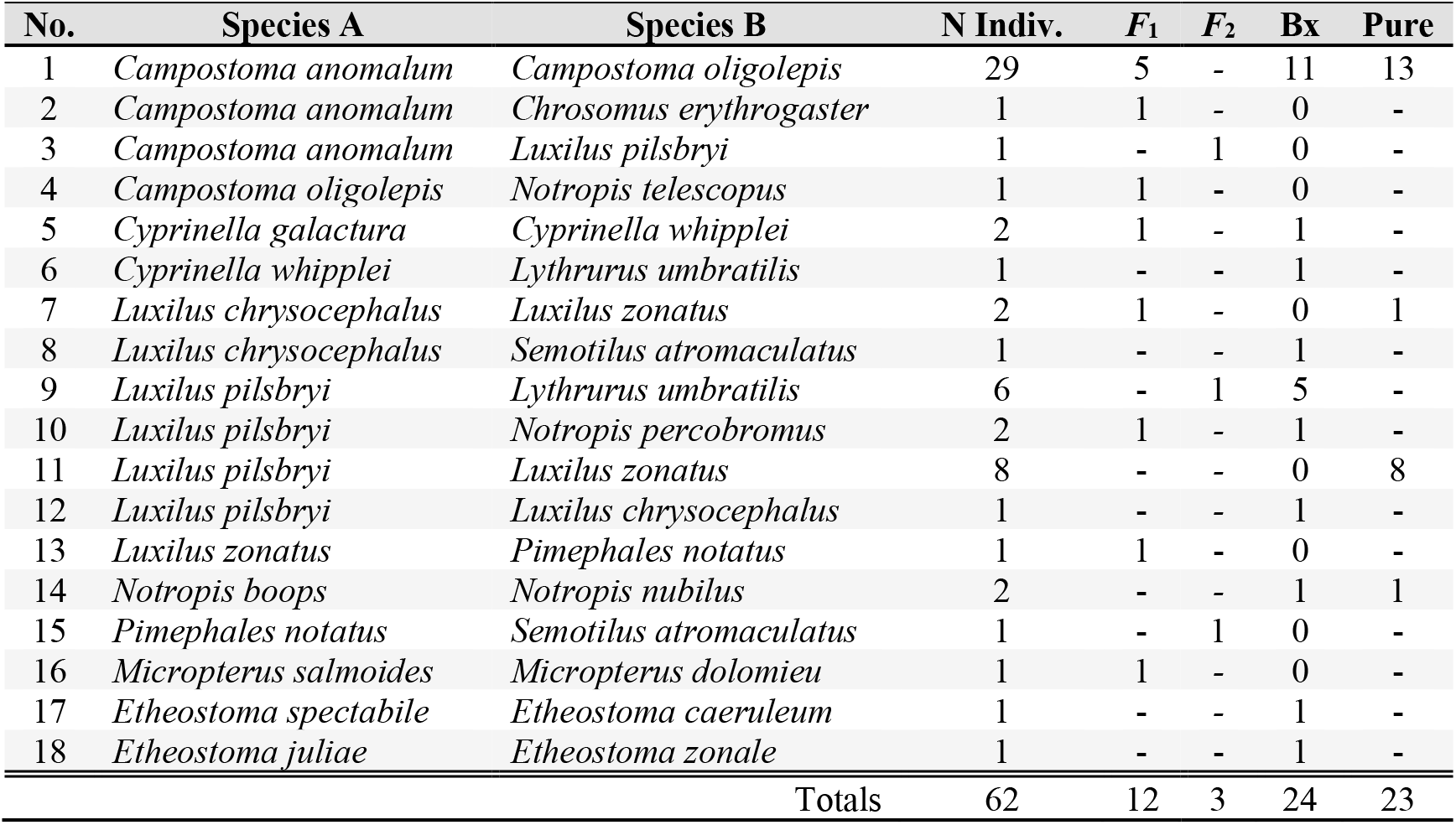
Observed genotypic frequency classes of hybrid individuals inferred from NewHybrids analysis for 18 hybridizing fish species-pairs collected across the White River basin, USA. Note that 8 multi-specific hybrids were not included. Putative hybrids were assigned to a genotype frequency class (*F*_1_, *F*_2_, Bx[=backcross], pure) based on Bayesian posterior probability >0.70.

### (c) Correlates of hybridization

Genetic differentiation among species (Table 4; Supplement S22) was significantly lower for hybridizing versus non-hybridizing pairs (mean *F*_ST_ = 0.86 vs. mean *F*_ST_ = 0.93; *W*=1521;*p*=0.002). The number of co-occurrences between hybridizing (mean=8.4) and non-hybridizing pairs (mean=7.1) did not differ significantly (*W*=960; *p*=0.76).

**TABLE 4.** Mean, minimum, and maximum values of Weir and Cockerham’s pairwise *F*_ST_ calculated among species within families collected across the White River Basin, USA. Families Cottidae and Fundulidae were represented by only two species each, hence one value of *F*_ST_.

| Family | Mean $F_{ST}$ | Min $F_{ST}$ | Max $F_{ST}$ |
| --- | --- | --- | --- |
| Centrarchidae | 0.94 | 0.85 | 0.98 |
| Cottidae | - | 0.82 | 0.82 |
| Fundulidae | - | 0.98 | 0.98 |
| Ictaluridae | 0.81 | 0.66 | 0.91 |
| Leuciscidae | 0.92 | 0.65 | 0.99 |
| Percidae | 0.93 | 0.78 | 0.98 |

At least one hybrid individual occurred within 42/75 communities (56%). We identified 12 environmental variables with a significant ability to reduce predictive error (i.e., VIMP) based on random permutation of predictors in the random forest (Supplement S23-S24). From these, just four were used to maximize the predictive capacity of the final model based on minimal depth selection (Figure 3), including species richness (VIMP=0.09), protected area extent (pac_pc_use; VIMP=0.04), mean precipitation in May (pre_mm_c05; VIMP=0.03), and mean annual precipitation (pre_mm_cyr; VIMP=0.03). The accuracy of the model was satisfactory: *G*-mean=0.73 and normalized Brier score=0.78. A similar misclassification error occurred for communities with and without hybrids (0.26 vs. 0.27, respectively).

**FIGURE 3.**
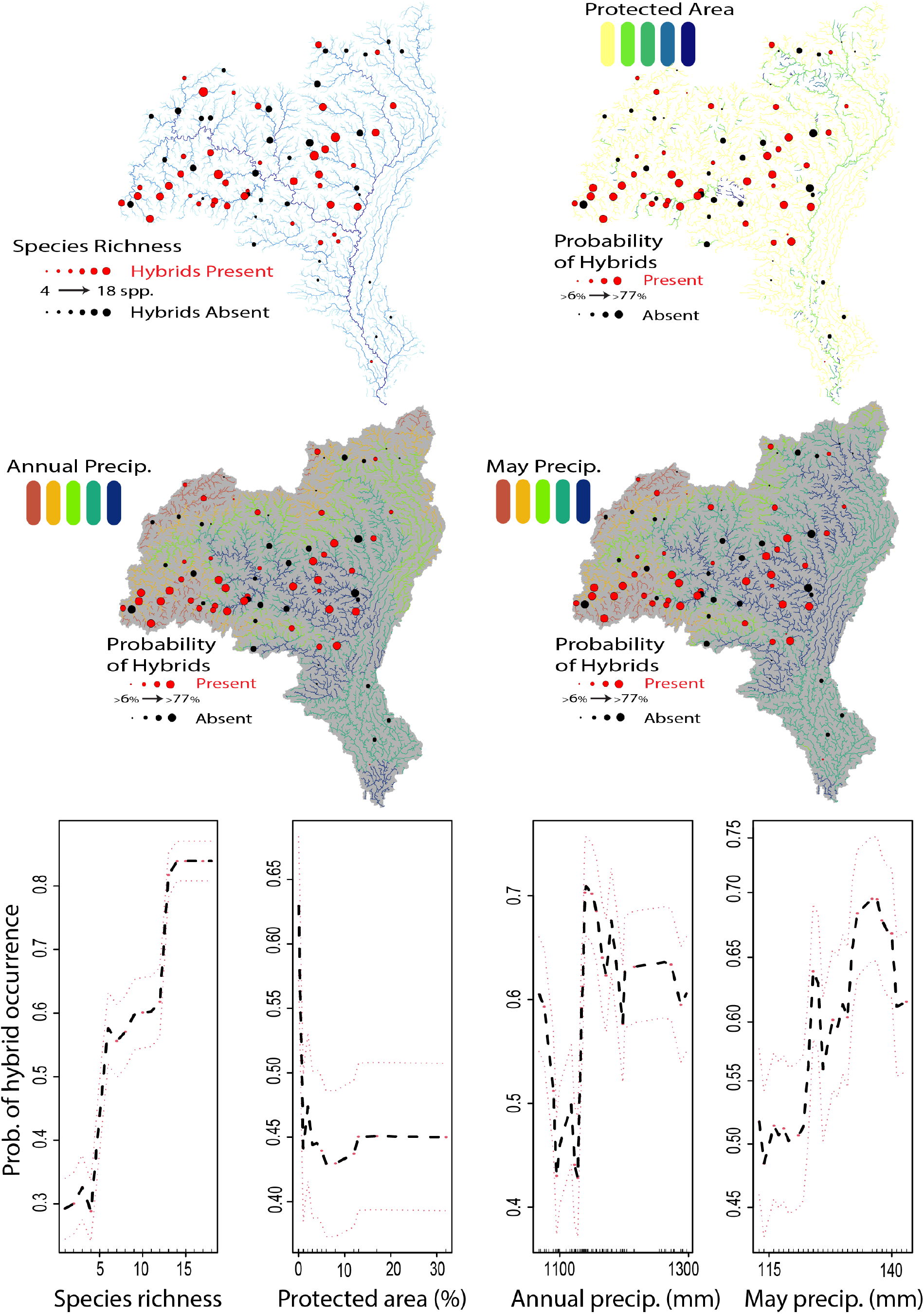
Significant predictors of hybridization across the White River Basin, USA. Maps depict collection sites with circles (red indicates presence of a hybrid). In the first map, site-diameter is scaled by species richness, while in the remaining maps, it is scaled by the probability of hybrid occurrence based on the model. Basin backgrounds differ for enhanced visibility. Below the maps are partial marginal effects plots showing relationships between probability of hybrid individuals occurring per four predictive variables. The mean (black) and 95% confidence intervals (red) are shown, along with distribution of variable values across sites (black tick marks above x-axis).

Neither species richness nor protect area extent were strongly correlated with other predictor variables (*r* >0.70; Pearson correlation). The correlation coefficient between mean May and annual precipitation was *r*=0.61. Both precipitation variables were strongly and positively correlated with other climatic variables related to temperature, evapotranspiration, and soil water content. Annual precipitation was also strongly but negatively correlated with river network position (distance from network outlet; DIST_DN_KM). Although not selected for the final model, eight additional variables were significantly associated with hybridization based on VIMP, including indices of human impact (*N*=2), road density (*N*=1), annual snow cover (*N*=1), and vegetative cover (*N*=4) (Supplement S23-24).

## 4. Discussion

Hybridization and introgression can impact fitness, facilitate gene exchange among species, or generate new lineages [5,6,55–57]. While hybridization has been widely documented at the per-species level, its occurrence within communities is expected to be rare; otherwise, species boundaries would seriously deteriorate [2]. Despite these expectations, hybrid prevalence at the community level is largely unknown, given that most studies to date focus on single species-pairs (or but a few closely related pairs) [55].

This study quantified hybridization at the community level without *a priori* assumptions of putative hybridization, thus serving as an appropriate broad-scale model without species-specific biases. It provided a more holistic survey of hybridization across the riverscape and thus stood in contrast to more conventional studies based on single species-pair evaluations.

### (a) Frequency of natural hybridization

The relevance of hybridization in ecology and evolution is underscored in our study by the detection of hybrids within four (of six) families (67%), involving 24 (of 33) species (73%). We documented hybridization among 18 unique species-pairs, 10 of which have been reported previously [20,58–62]. Hybridization across our study region was overwhelmingly within leuciscids, a family oft-dominating within Ozark stream communities [63]. The unbalanced number of representatives from each family reflects that and should not necessarily be interpreted as differences in the propensity to hybridize among the families more generally. Surprisingly, few hybrids were detected within Centrarchidae and Percidae, despite an extensively-documented presence [19,64,65].

Hybridization was encountered more frequently than anticipated, whether viewed per-individual (2.4%) or per-species (73%). Previous estimates in plants and animals are lower than found herein (0.002–0.06%) [15,66,67]. If rates among our study species were comparable to the previous (e.g., 0.1%), we would have expected but 2–3 hybrids. Earlier estimates relied on morphological identification, a methodology less sensitive in detecting later-generation hybrids [19,62]. Additionally, rates herein also reflect both our breadth of individuals evaluated, as well as potential comparisons so tested. While our study is but a subsample of individuals from the region and lacks representatives from every extant species, we nevertheless attempted to reduce unsampled lineages that could bias inferences and allow undetected or misclassified hybrids [68,69].

Hybridization is seemingly greater in fishes than in other vertebrates due to their external fertilization within aqueous environs [1,19,70]. Similar per-individual rates (0–4%) were identified in marine fishes [71], with even higher rates (22.5%) among invasive Mississippi River carp [72]. Published per-species estimates varied from 1–10% across animals and ~25% for plants [2,73]. Although differences in the former are apparent from literature and museum records, one could ask if it reflects an actual biological signal or variance in diversity and/or research effort. For example, once adjustments were made for diversity and research intensity, rates were mostly homogenous among taxonomic groups compiled in a meta-analysis, but with the caveat that rates for fishes were still demonstrably higher than expected [18].

Our per-species rate was highest within our most specious group (Leuciscidae; Table 2). This parallels previous meta-analyses identifying disproportionately high hybridization rates within minnows [1,19,20]. Interestingly, leuciscid breeding behaviors—especially nest building and association— significantly predicted hybridization rates across the clade [20]. Many minnows broadcast gametes widely or affix them onto substrates, often shared among species [20,74], with as many as six simultaneously employing the same gravel substrate at once [1]. In addition, minnows comprise the most diverse and widely distributed North American stream fish family and numerically dominate stream communities [75]. They thus encounter broad environmental heterogeneity, demonstrate a high degree of sympatry, and exhibit uneven species abundances, which promote hybridization [19].

### (b) Introgression

The evidence of backcrossing between hybrids and parentals suggests that hybrids can be viable and fertile, thus facilitating introgression [67]. This genetic exchange can provide a source of novelty for evolutionary forces to act upon [73], and the prevalence found here provides some indication as to how probable an extraneous genetic contribution can be. We identified 24 individuals seemingly resulting from parental/hybrid backcrosses spanning ten species-pairs and two families (Leuciscidae, Percidae). It is intriguing that the most diverse group of North American freshwater fish—leuciscids—has an established reputation for frequent hybridization and is also a prolific contributor to gene exchange among species in our study. Introgressive hybridization has played a crucial role in cyprinid evolutionary history [67], and the evidence gathered here suggests it may continue to do so.

Questions remain regarding whether the introgression seen in our region is evolutionarily adaptive, maladaptive, or neutral. This may ultimately hinge upon differences among species and environments [76]. Answers would facilitate our capacity to predict and mitigate adverse outcomes of hybridization [77]. Climate-mediated environmental shifts will predictably exacerbate hybridization, especially for fishes whose life histories are sensitive to spawning temperatures and streamflow [12]. Yet we recognize that hybridization is not necessarily negative. For example, adaptive introgression between generalist and specialist rainbow fishes has seemingly diminished climate change vulnerability in admixed individuals compared to pure populations [10]. It is thus viewed as a valuable component of “evolutionary rescue” and an underappreciated conservation tool [10].

### (c) Correlates of hybridization

Our data concur with the premise that hybridization is strongly influenced by both divergences among species and also environmental factors. We focus our discussion on the latter, per the novelty of our data and the previously established support for the former [20–22]. In synopsis, communities were more likely to harbor hybrids when: (i) Greater species richness existed; (ii) Protected area within the catchment was limited; and (iii) Habitats were prone to more precipitation.

While hybridizing and non-hybridizing species-pairs did not differ significantly in their co-occurrence, we did note that communities where more species occur together (greater richness) also possessed more hybrids (Figure 3). Although seemingly intuitive, the opposite is suggested at larger spatial and temporal scales where species-richness diminishes with greater niche availability at biogeographic borders [16,23]. Empirical results are few, and conclusions varied: For example, the number of plant hybrids across US counties was significantly related to species richness [17]. However, hybridization and species richness were not significantly related among coral reefs [14,16].

The extent of protected areas within the total watershed upstream of the reach was negatively associated with basin-level hybridization (Figure 3). Similarly, anthropogenic impacts were positively associated with hybridization (Supplement S24). Environmental perturbations in general, and specifically those anthropogenic, loom large in the hybridization literature [19,28]. Stable, more pristine environments are expected to harbor fewer hybrids than those perturbed due mainly to a breakdown of reproductive isolating mechanisms facilitated by translocations, habitat modifications, and ongoing climate change [28,78].

Greater mean precipitation within the reach catchment (annually and in May) was also associated with elevated hybridization (Figure 3). More precipitation is associated with warmer and more downstream communities in our system (i.e., more flow). Elevated levels of hybridization in these communities are potentially driven by flooding magnitude, particularly given the combination of higher precipitation and lower network position. Horton stream order and catchment size promote the magnitude of flooding [79], as compounded by the precipitation gradient of our study system [80]. Additionally, the Hydrologic Disturbance Index (a compendium of several anthropogenic impacts) is more significant in streams with larger drainage areas and lower gradients (i.e., further downstream), which in turn has been shown to promote variance in fish community composition [31].

Flooding can affect fish spawning in several ways: promotion of spawning activity [80]; concentration of fishes within refugia [53,81]; disturbance of spawning habitats and nests [82]; displacement of oviposited eggs [53]; and elevation of discharge/turbidity, which weakens those sensory cues (e.g., visual, olfactory, environmental) that sustain breeding isolation [25–27]. The inclusion of May precipitation in our most predictive models supports the above, in that study species (save two cottids) spawn in May/June [83].

## 5. Conclusion

Hybridization occurs more frequently than expected in the White River Basin and is predictable based on specifics of the environment. Although recognized as a creative evolutionary force, it is also considered a maladaptive threat. Further research may blueprint an even more complicated future and potentially demonstrate that groups (such as minnows) thrive proportional to their prolific gene exchange. Moreover, hybridization is predicted to increase in frequency with global environmental change [11,12], a recognition consistent with our finding of hybrid occurrence in lockstep with climate-related variables. Therefore, baseline estimates are required to gauge the increase in hybridization, predict which ecosystems will be so impacted (and how severely), and promote a more robust conservation/management strategy that allows those impacts to be understood and adjudicated (if so needed) [10]. Future studies like ours will be performed at the whole-genome level, with a resolution more robust for detecting and untangling hybridization and its genomic consequences [24].

## Acknowledgments

We thank M. Flurry, M. George, T. Goodhart, K. Hollar, and M. Reed, who assisted with DNA extractions. The Arkansas High-Performance Computing Center provided analytical resources.

## Author contributions

ZDZ conceived the research with input from all authors. Specimen collection was done by ZDZ & TKC. Laboratory work, bioinformatics, data analysis, and manuscript drafting were done by ZDZ. All authors contributed to the analysis interpretation and critically revising the manuscript. MRD and MED administered funding through their endowments.

## Data availability statement

Raw sequence files are accessioned in the NCBI GenBank Sequence Read Archive (SRA) BioProject: PRJNA809538. [84]. SNP alignments and R code are archived on Open Science Framework [85]

