## Supplemental Material for "A community genomics approach to natural hybridization"

**TABLE OF CONTENTS**

| Supplement S1 | Supplemental Methods | p. 2 |
| --- | --- | --- |
| Supplement S2 | Power analysis: *Micropterus dolomieu* X *Micropterus salmoides* | p. 3 |
| Supplement S3 | Power analysis: *Etheostoma caeruleum* X *Etheostoma spectabile* | p. 4 |
| Supplement S4 | Power analysis: *Etheostoma juliae* X *Etheostoma zonale* | p. 5 |
| Supplement S5 | Power analysis: *Campostoma anomalum* X *Campostoma oligolepis* | p. 6 |
| Supplement S6 | Power analysis: *Campostoma anomalum* X *Chrosomus erythrogaster* | p. 7 |
| Supplement S7 | Power analysis: *Campostoma anomalum* X *Luxilus pilsbryi* | p. 8 |
| Supplement S8 | Power analysis: *Campostoma oligolepis* X *Notropis telescopus* | p. 9 |
| Supplement S9 | Power analysis: *Cyprinella whipplei* X *Cyprinella galactura* | p. 10 |
| Supplement S10 | Power analysis: *Cyprinella whipplei* X *Lythrurus umbratilis* | p. 11 |
| Supplement S11 | Power analysis: *Luxilus chrysocephalus* X *Luxilus zonatus* | p. 12 |
| Supplement S12 | Power analysis: *Luxilus chrysocephalus* X *Semotilus atromaculatus* | p. 13 |
| Supplement S13 | Power analysis: *Luxilus pilsbryi* X *Luxilus chrysocephalus* | p. 14 |
| Supplement S14 | Power analysis: *Luxilus pilsbryi* X *Luxilus zonatus* | p. 15 |
| Supplement S15 | Power analysis: *Luxilus pilsbryi* X *Lythrurus umbratilis* | p. 16 |
| Supplement S16 | Power analysis: *Luxilus pilsbryi* X *Notropis percobromus* | p. 17 |
| Supplement S17 | Power analysis: *Luxilus zonatus* X *Pimephales notatus* | p. 18 |
| Supplement S18 | Power analysis: *Notropis boops* X *Notropis nubilus* | p. 19 |
| Supplement S19 | Power analysis: *Pimephales notatus* X *Semotilus atromaculatus* | p. 20 |
| Supplement S20 | List of 18 hybridizing species pairs with literature references | p. 21 |
| Supplement S21 | Summary of *N*=70 hybrid individuals | p. 22 |
| Supplement S22 | Fst and co-occurrence for 137 Unique Intra-familial pairs | p. 24 |
| Supplement S23 | Significant environmental factors for predicting hybrid incidence via random forest | p. 26 |
| Supplement S24 | Marginal effects plots of significant environmental factors | p. 27 |
| Supplement S25 | Species codes used in the individual summary table and throughout code | p. 28 |

**SUPPLEMENT S1** Supplementary materials and methods

Collecting permits

Collecting permits were provided by: Arkansas Game & Fish Commission (#020120191); Missouri Dept. Wildlife Conservation (#18136); and US National Parks Service (NPS: Buffalo River Permit; BUFF-2017-SCI-0013).

Library preparation

Standardized DNA concentrations (1000 ng) were digested at 37°C with high-fidelity restriction enzymes *MspI* (5’-CCGG-3') and *PstI* (5’-CTGCAG-3') (New England Biosciences), bead-purified (Ampure XP; Beckman-Coulter Inc.), standardized to 100 ng, and then ligated with custom adapters containing in-line identifying barcodes (T4 Ligase; New England Biosciences). Individual samples were pooled in sets of 48 and size-selected from 326 to 426 bp (Pippin Prep; Sage Sciences). Illumina adapters and i7 index were added via 12-cycle PCR with Phusion high-fidelity DNA polymerase (New England Biosciences). A set of three libraries (3x48=144 individuals/lane) were pooled per-lane and sequenced single-end on the Illumina HiSeq 4000 platform (1x100bp; Genomics & Cell Characterization Core Facility; University of Oregon/ Eugene). Quality control checks, to include fragment analysis and quantitative real-time PCR, were performed at the core facility prior to sequencing.

Alignment and assembly free (AAF) phylogenies

Data assembly was performed in multiple stages: Pre-processing; Species validation and screening; and family-level alignment. Reads were first demultiplexed in ipyrad v.0.9.62 allowing up to one barcode mismatch, yielding individual FASTQ files containing raw reads (*N*=3,101). Individuals averaged >2 million reads, with those extremely low $(< \bar{x}-2\sigma)$ being removed (*N=59*). Species identifications were validated using an alignment and assembly-free (AAF) method (PHYLORAD). This method is comparable to alignment-based methods but with lower computational burdens, as accomplished by computing a pairwise distance matrix as the proportion of DNA substrings of length *k* (=*k*-mer) as a function of total unique identified *k*-mers, followed by phylogenetic reconstruction following the Fitch-Margoliash method. The same *k*-mer length was employed for read selection (*k*s) and reconstruction (*k*) = 21, with a k-mer filtering threshold (*n*) = 2. Phylogenies were inspected manually, with putative mis-identified individuals not clustering with the appropriate species-clade. These individuals were re-evaluated via vouchered specimens (*N*=249; 8%), and a assinged a different species ID due to misidentification.

Data processing and assembly

Individuals (*N*=3,042) were subsequently partitioned by family (*N*=6) and processed *de novo* in ipyrad to generate family-level assemblies. Adapters and primers were removed and reads with >5 low-quality bases (Phred<20) discarded. Clusters were assembled using an 85% identity threshold, with loci subsequently removed via conditional criteria to ensure high-quality data: <20x and >500x coverage per individual; >5% of consensus nucleotides ambiguous; >20% of nucleotides polymorphic; >8 indels present; or presence in <15% of individuals. Putative paralogs were removed if clusters displayed >2 alleles per site in consensus sequence or excessive heterozygosity (>5% consensus bases or >50% heterozygosity/site). Biallelic SNPs were further filtered in radiator, and removed if: Monomorphic; minor allele frequency <3%; mean coverage <20 or >200; missing data >30%; SNP position on read >91; and if HWE lacking in one or more species (α=0.0001). Only one SNP was retained per locus (i.e., that which maximized minor allele count). Finally, individuals having >50% missing data were excluded.

**SUPPLEMENT S2** Power analysis for detecting genotype frequency classes via HYBRIDDETECTIVE workflow. Results are shown here for *Micropterus dolomieu* X *Micropterus salmoides*. P1 and P2 = pure; F1 and F2 = first filial and second filial hybrids; BC1 and BC2 = backcrosses.

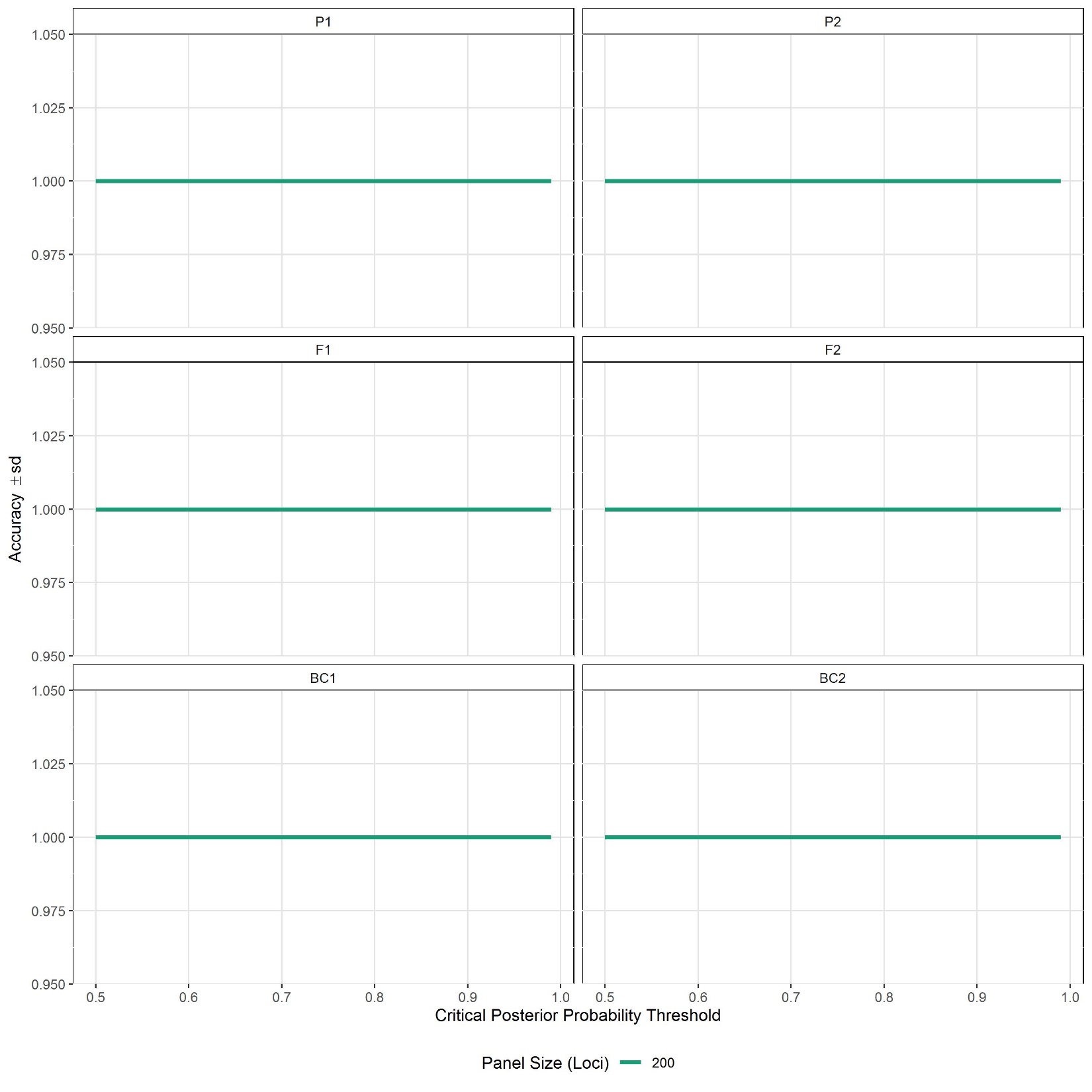

**SUPPLEMENT S3** Power analysis for detecting genotype frequency classes via HYBRIDDETECTIVE workflow. Results are shown here for *Etheostoma caeruleum* X *Etheostoma spectabile*. P1 and P2 = pure; F1 and F2 = first filial and second filial hybrids; BC1 and BC2 = backcrosses.

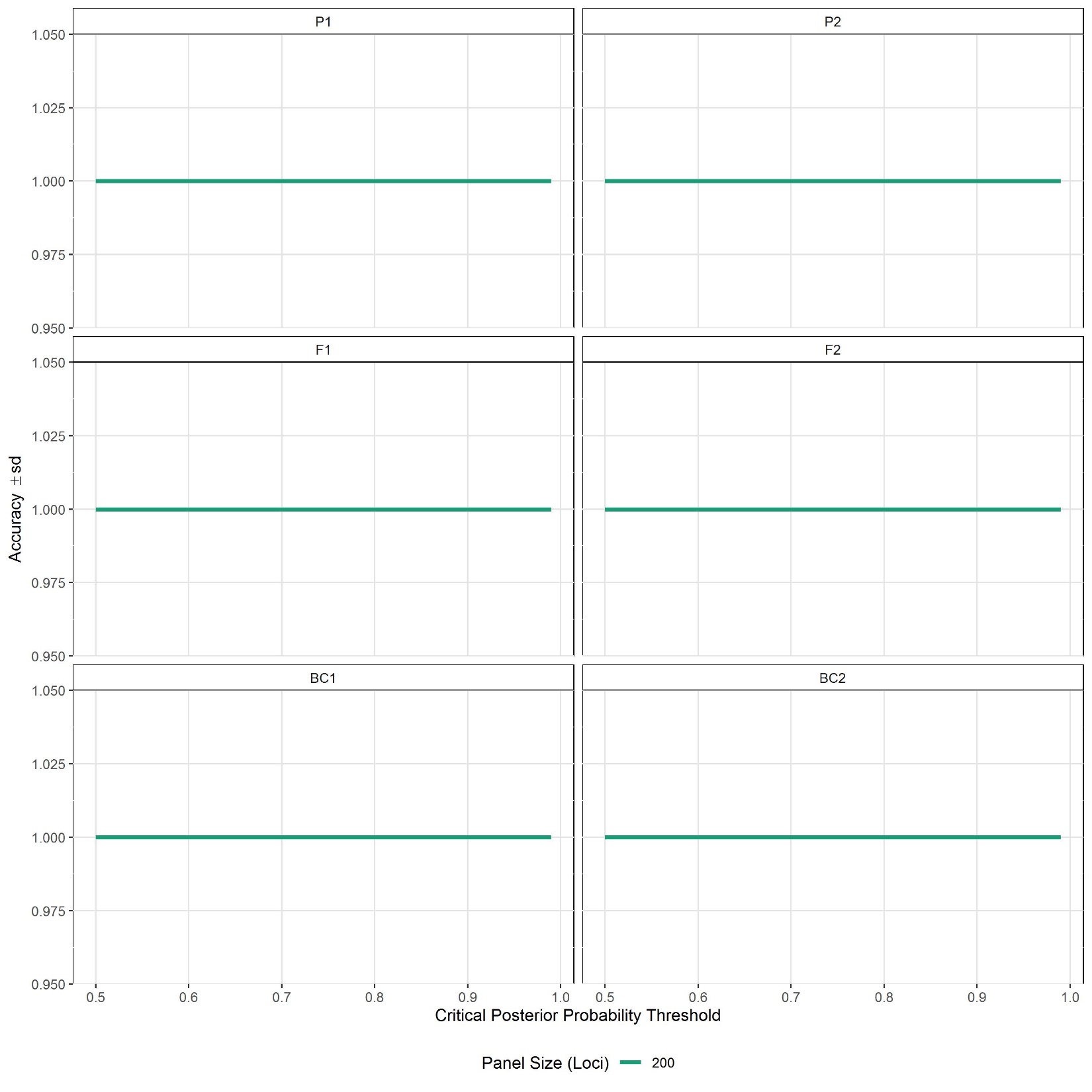

**SUPPLEMENT S4** Power analysis for detecting genotype frequency classes via HYBRIDDETECTIVE workflow. Results are shown here for *Etheostoma juliae* X *Etheostoma zonale*. P1 and P2 = pure; F1 and F2 = first filial and second filial hybrids; BC1 and BC2 = backcrosses.

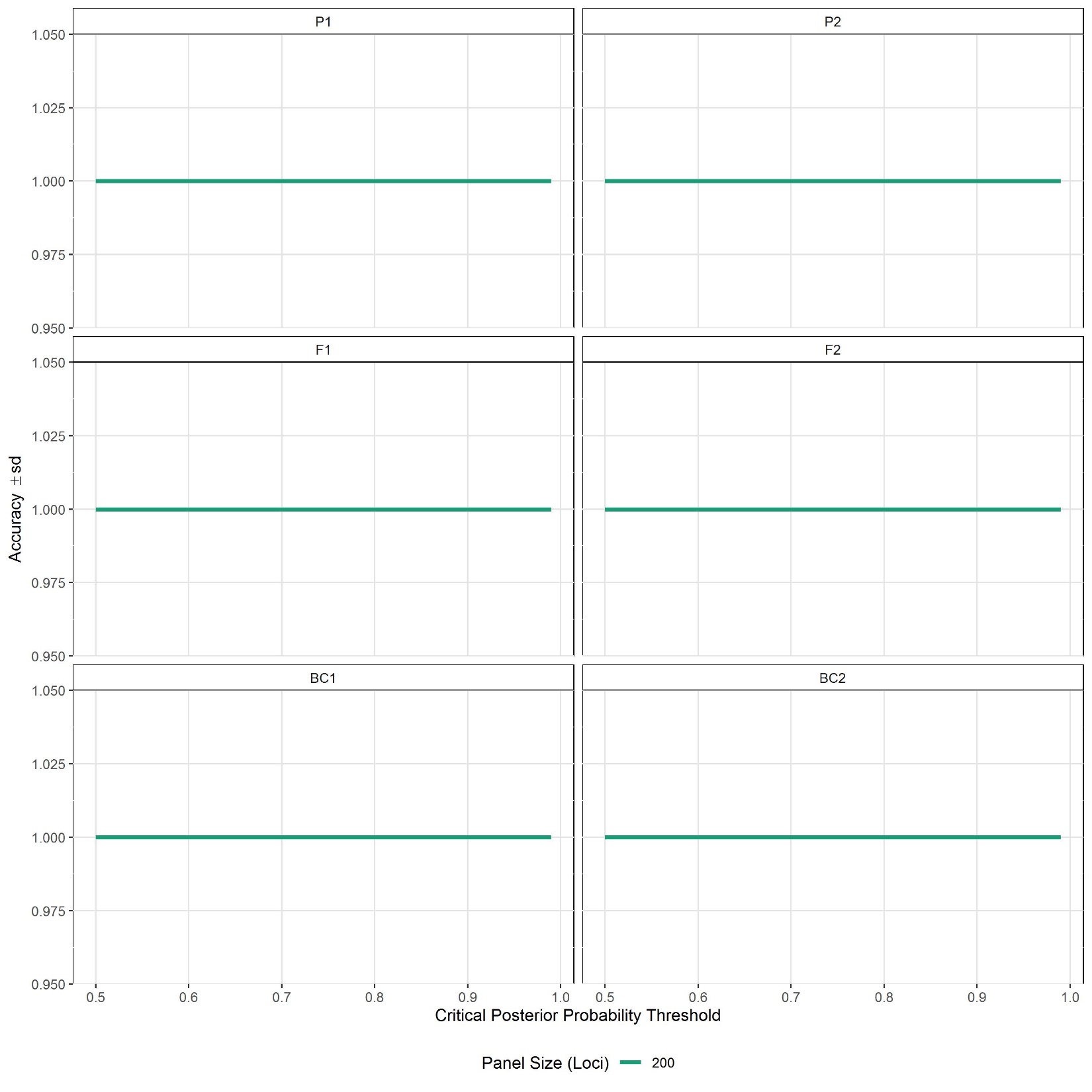

**SUPPLEMENT S5** Power analysis for detecting genotype frequency classes via HYBRIDDETECTIVE workflow. Results are shown here for *Campostoma anomalum* X *Campostoma oligolepis*. P1 and P2 = pure; F1 and F2 = first filial and second filial hybrids; BC1 and BC2 = backcrosses.

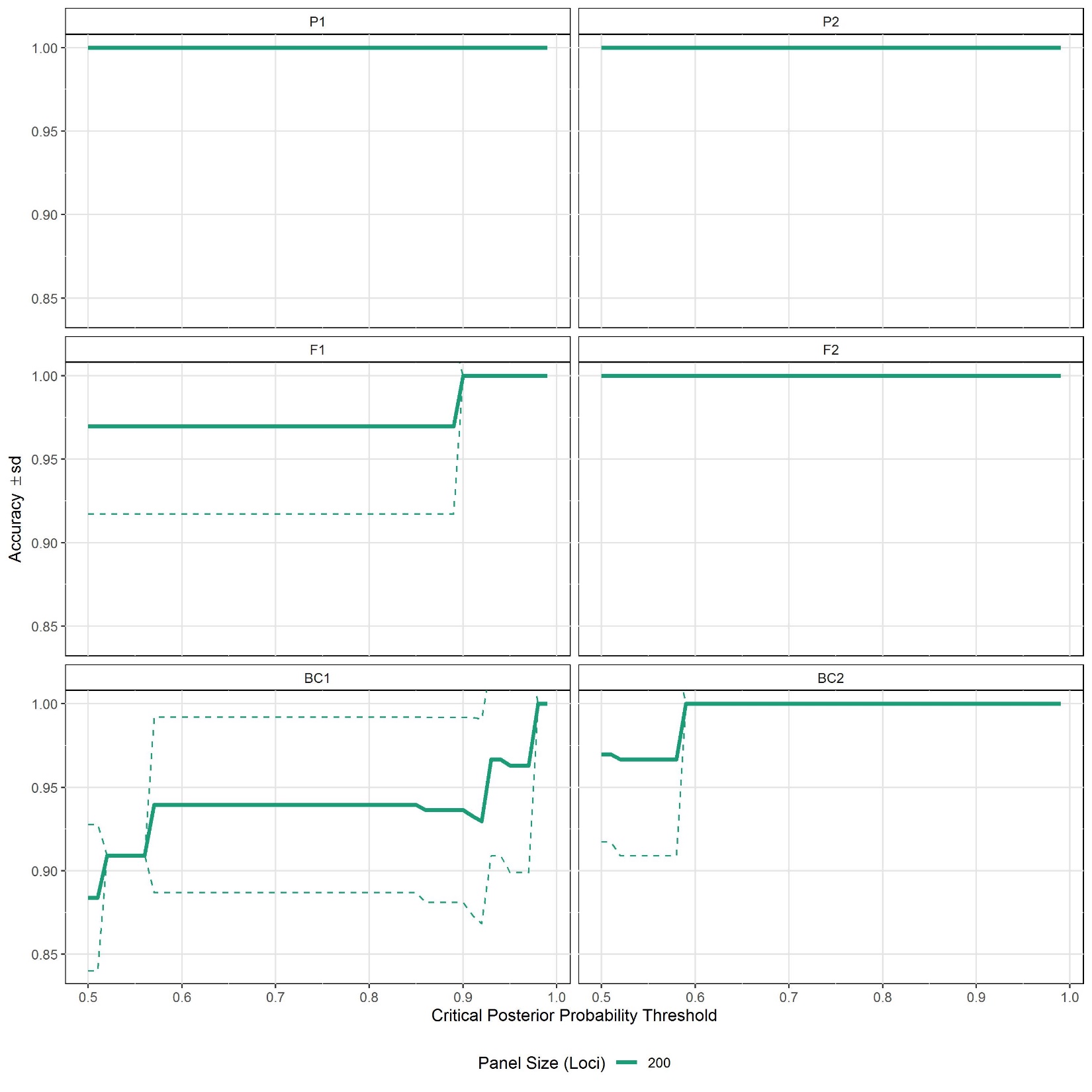

**SUPPLEMENT S6** Power analysis for detecting genotype frequency classes via HYBRIDDETECTIVE workflow. Results are shown here for *Campostoma anomalum* X *Chrosomus erythrogaster*. P1 and P2 = pure; F1 and F2 = first filial and second filial hybrids; BC1 and BC2 = backcrosses.

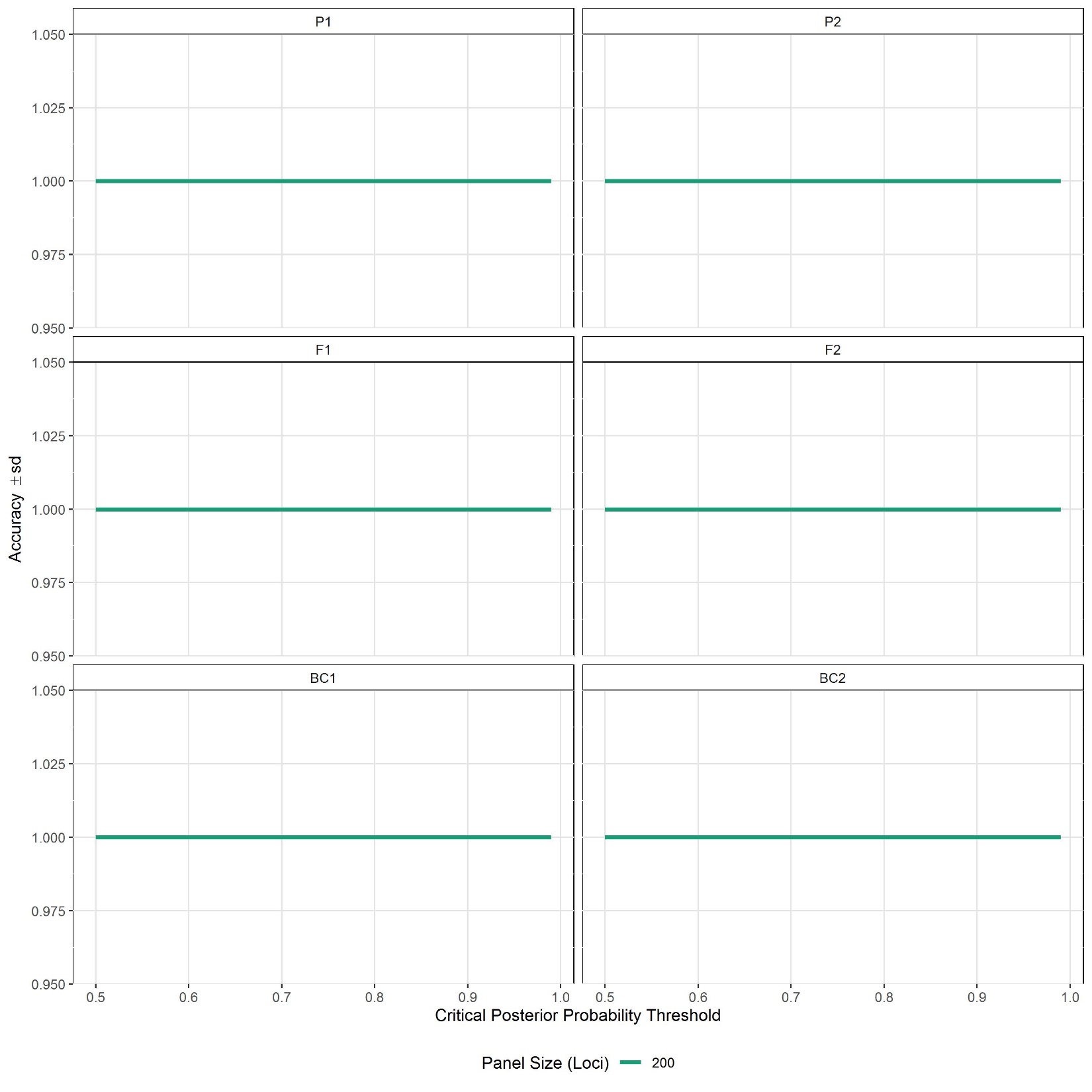

**SUPPLEMENT S7** Power analysis for detecting genotype frequency classes via HYBRIDDETECTIVE workflow. Results are shown here for *Campostoma anomalum* X *Luxilus pilsbryi*. P1 and P2 = pure; F1 and F2 = first filial and second filial hybrids; BC1 and BC2 = backcrosses.

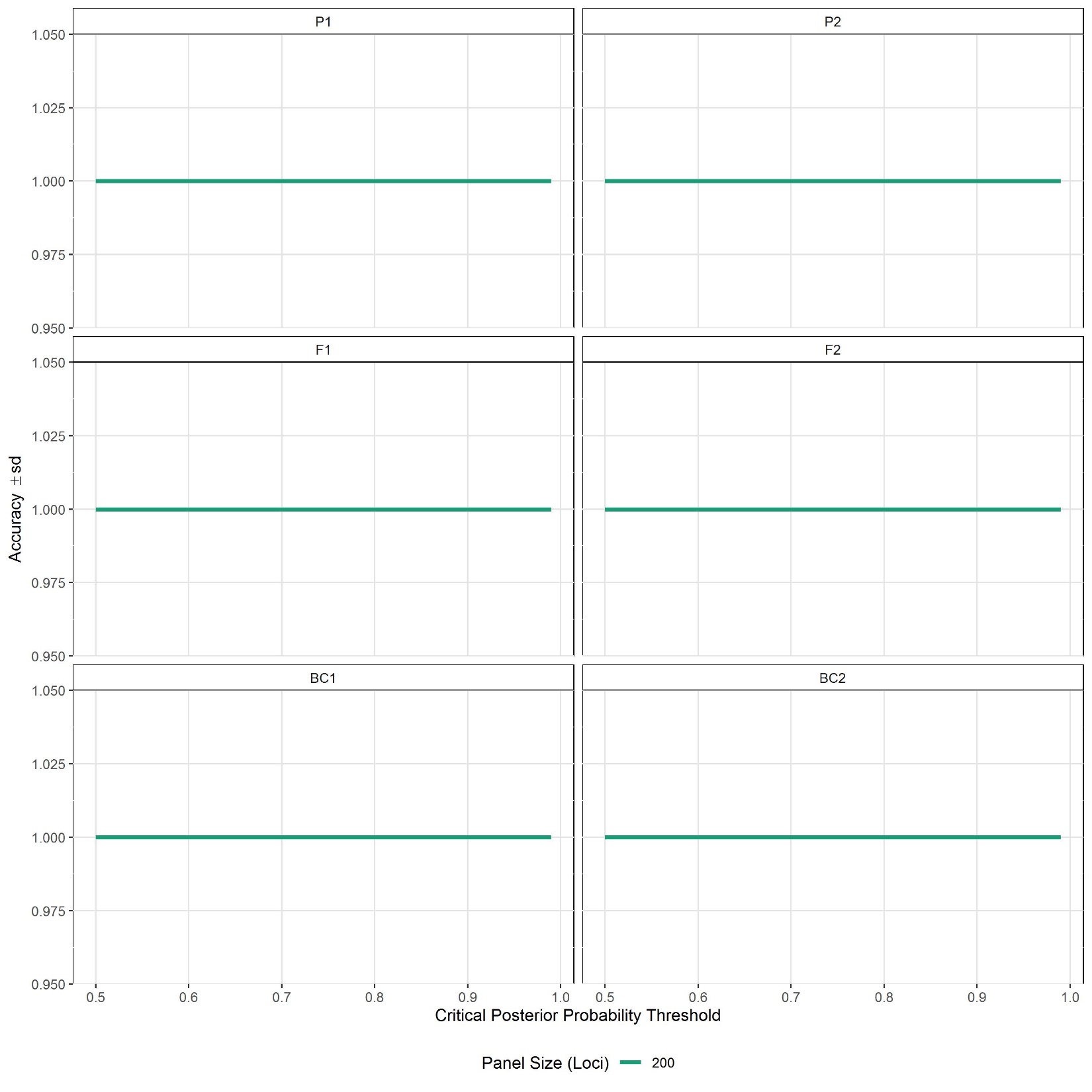

**SUPPLEMENT S8** Power analysis for detecting genotype frequency classes via HYBRIDDETECTIVE workflow. Results are shown here for *Campostoma oligolepis* X *Notropis telescopus*. P1 and P2 = pure; F1 and F2 = first filial and second filial hybrids; BC1 and BC2 = backcrosses.

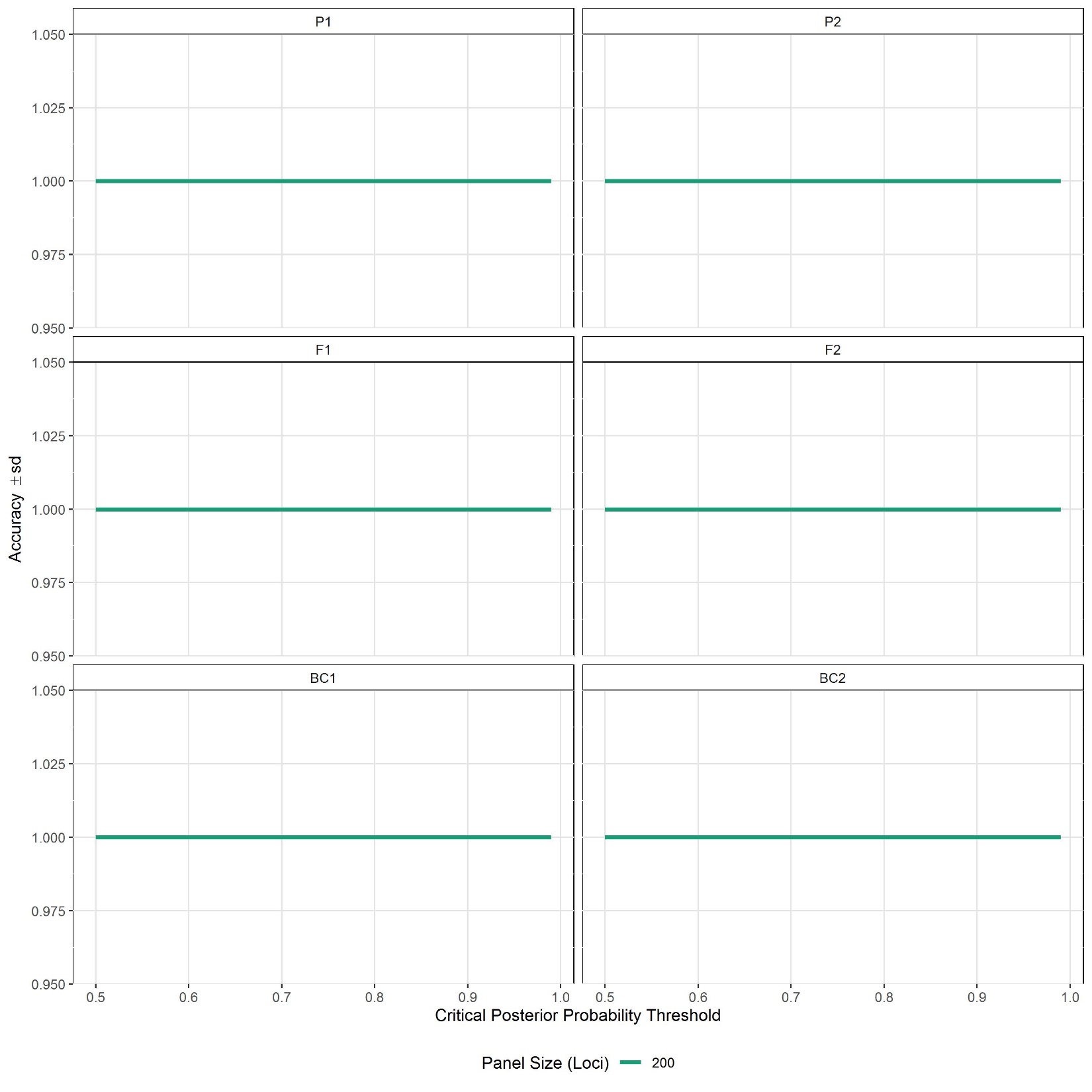

**SUPPLEMENT S9** Power analysis for detecting genotype frequency classes via HYBRIDDETECTIVE workflow. Results are shown here for *Cyprinella whipplei* X *Cyprinella galactura*. P1 and P2 = pure; F1 and F2 = first filial and second filial hybrids; BC1 and BC2 = backcrosses.

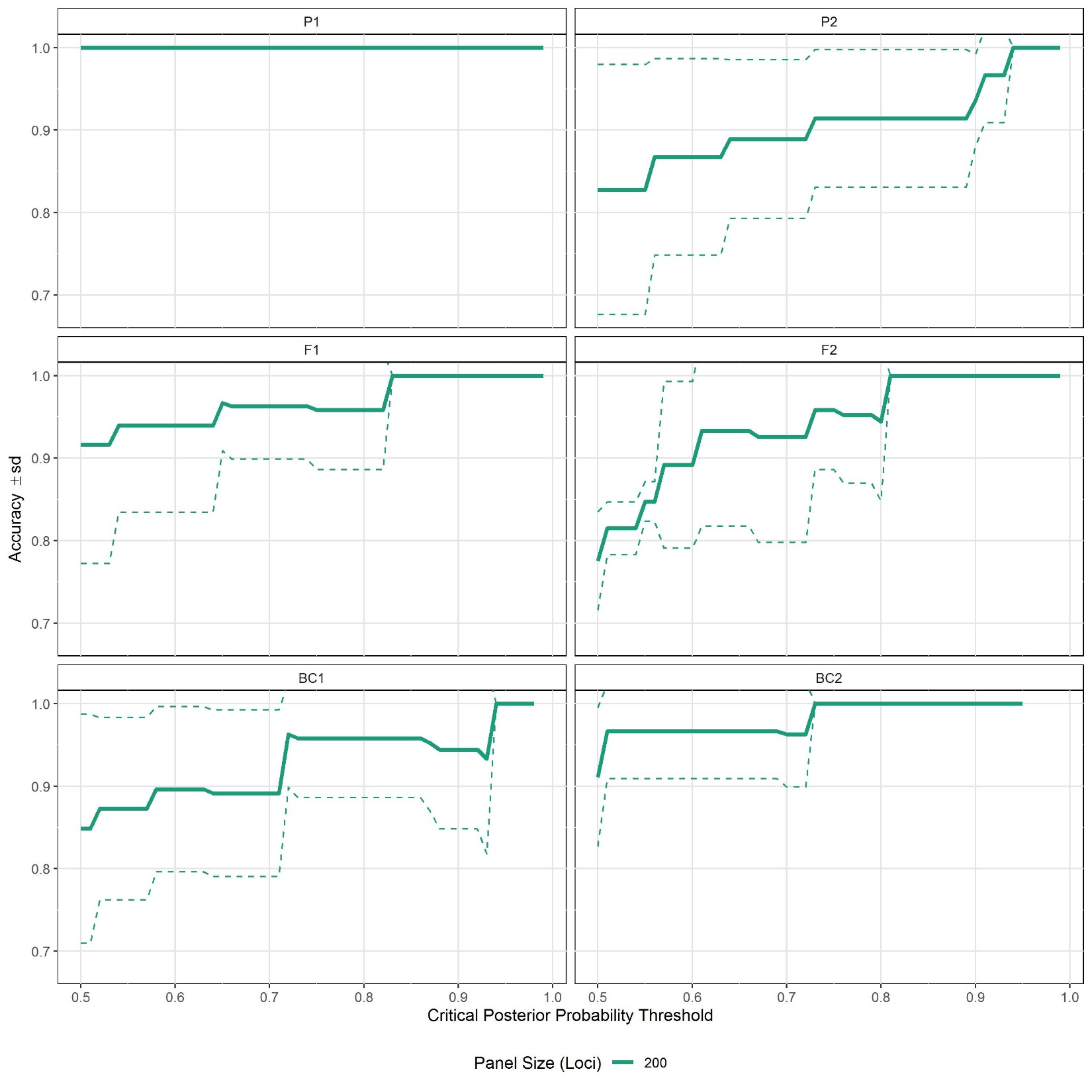

**SUPPLEMENT S10** Power analysis for detecting genotype frequency classes via HYBRIDDETECTIVE workflow. Results are shown here for *Cyprinella whipplei* X *Lythrurus umbratilis*. P1 and P2 = pure; F1 and F2 = first filial and second filial hybrids; BC1 and BC2 = backcrosses.

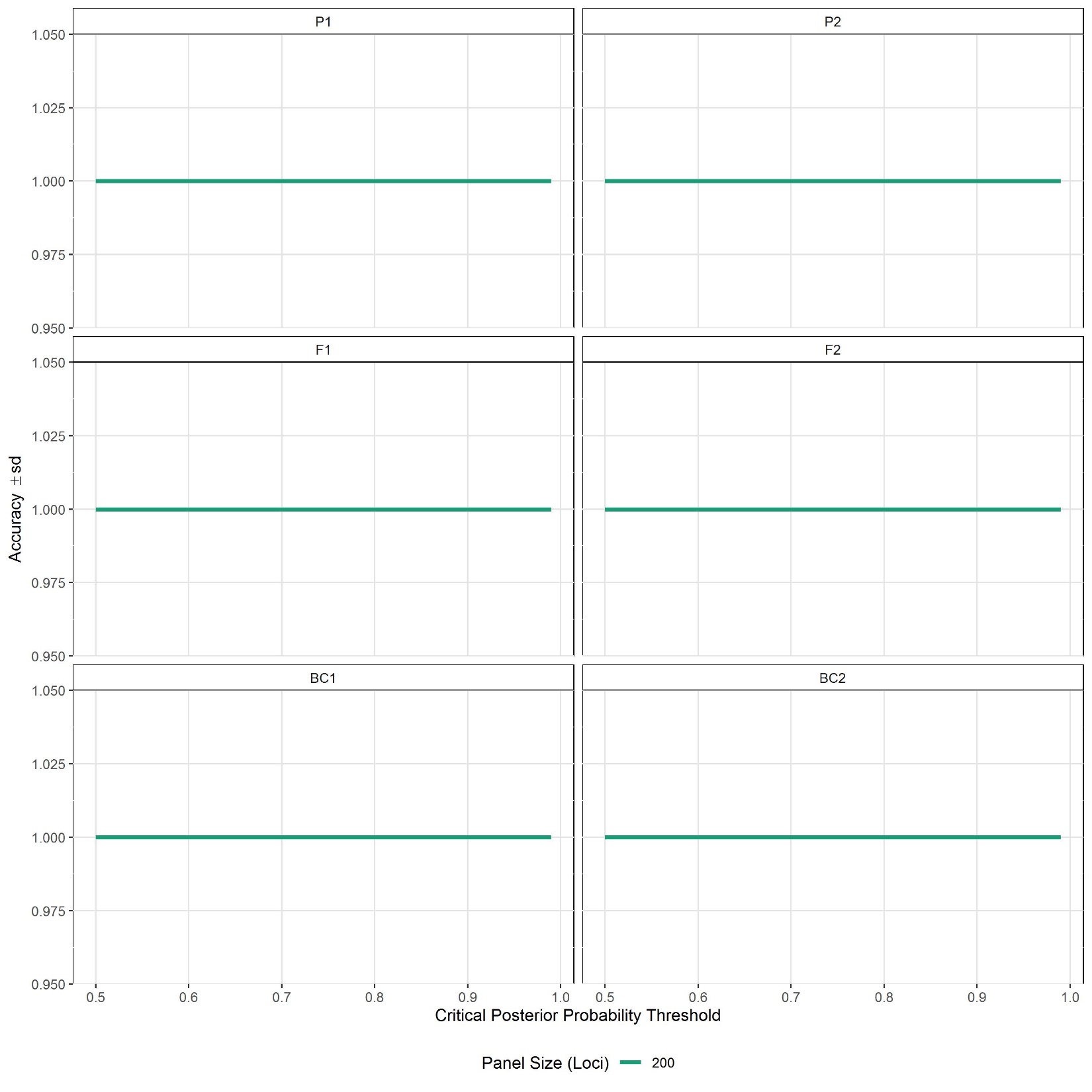

**SUPPLEMENT S11** Power analysis for detecting genotype frequency classes via HYBRIDDETECTIVE workflow. Results are shown here for *Luxilus chrysocephalus* X *Luxilus zonatus*. P1 and P2 = pure; F1 and F2 = first filial and second filial hybrids; BC1 and BC2 = backcrosses.

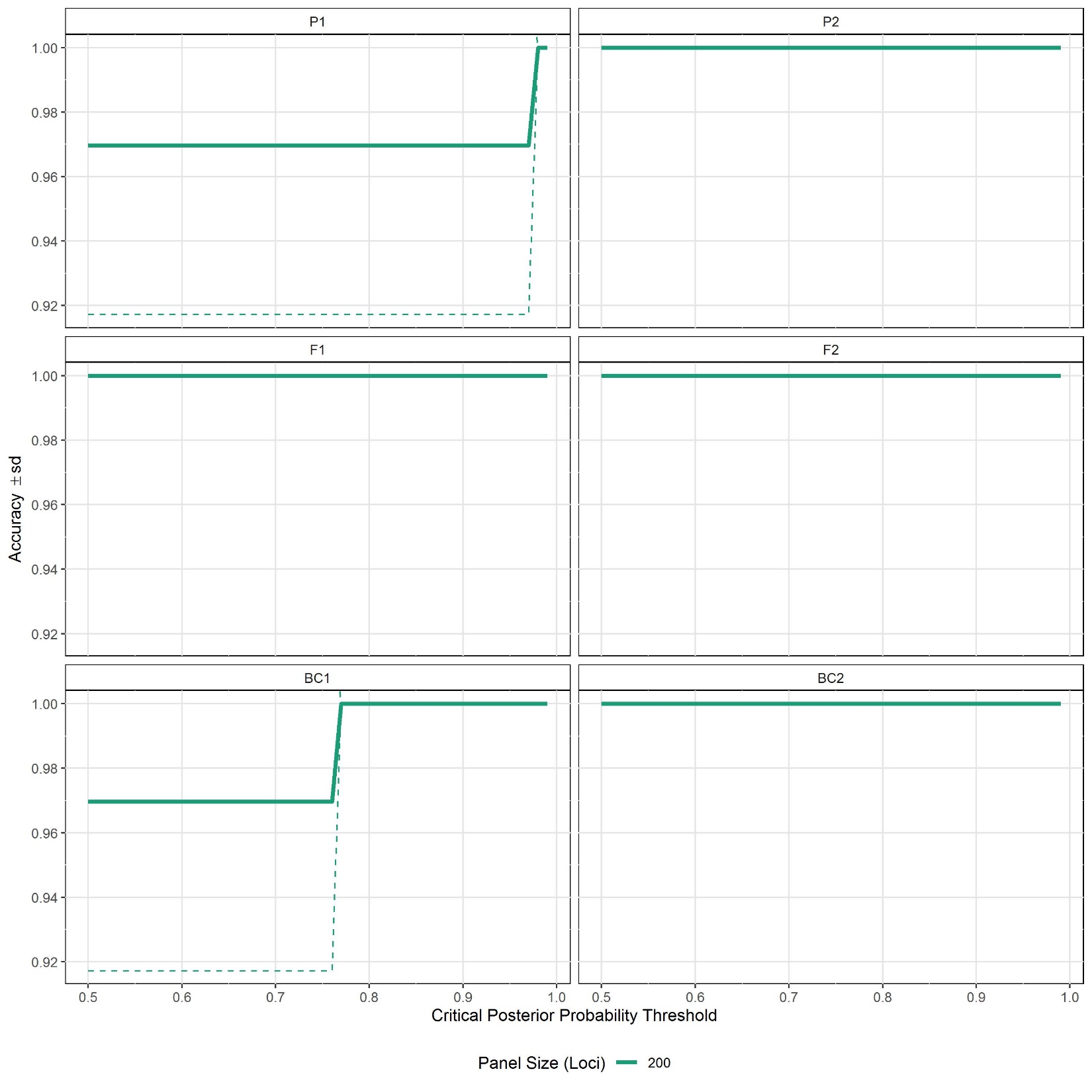

**SUPPLEMENT S12** Power analysis for detecting genotype frequency classes via HYBRIDDETECTIVE workflow. Results are shown here for *Luxilus chrysocepalus* X *Semotilus atromaculatus*. P1 and P2 = pure; F1 and F2 = first filial and second filial hybrids; BC1 and BC2 = backcrosses.

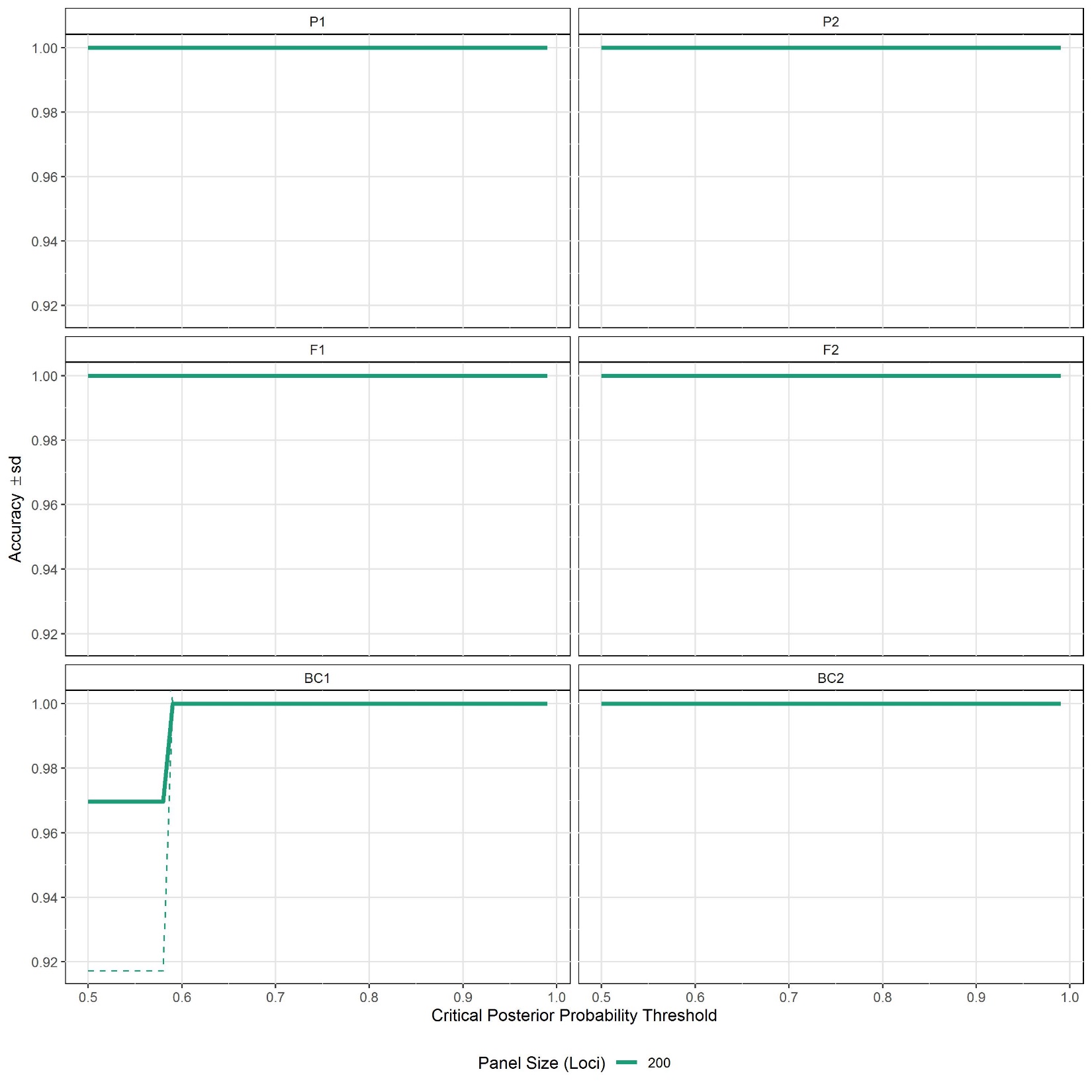

**SUPPLEMENT S13** Power analysis for detecting genotype frequency classes via HYBRIDDETECTIVE workflow. Results are shown here for *Luxilus pilsbryi* X *Luxilus chrysocephalus*. P1 and P2 = pure; F1 and F2 = first filial and second filial hybrids; BC1 and BC2 = backcrosses.

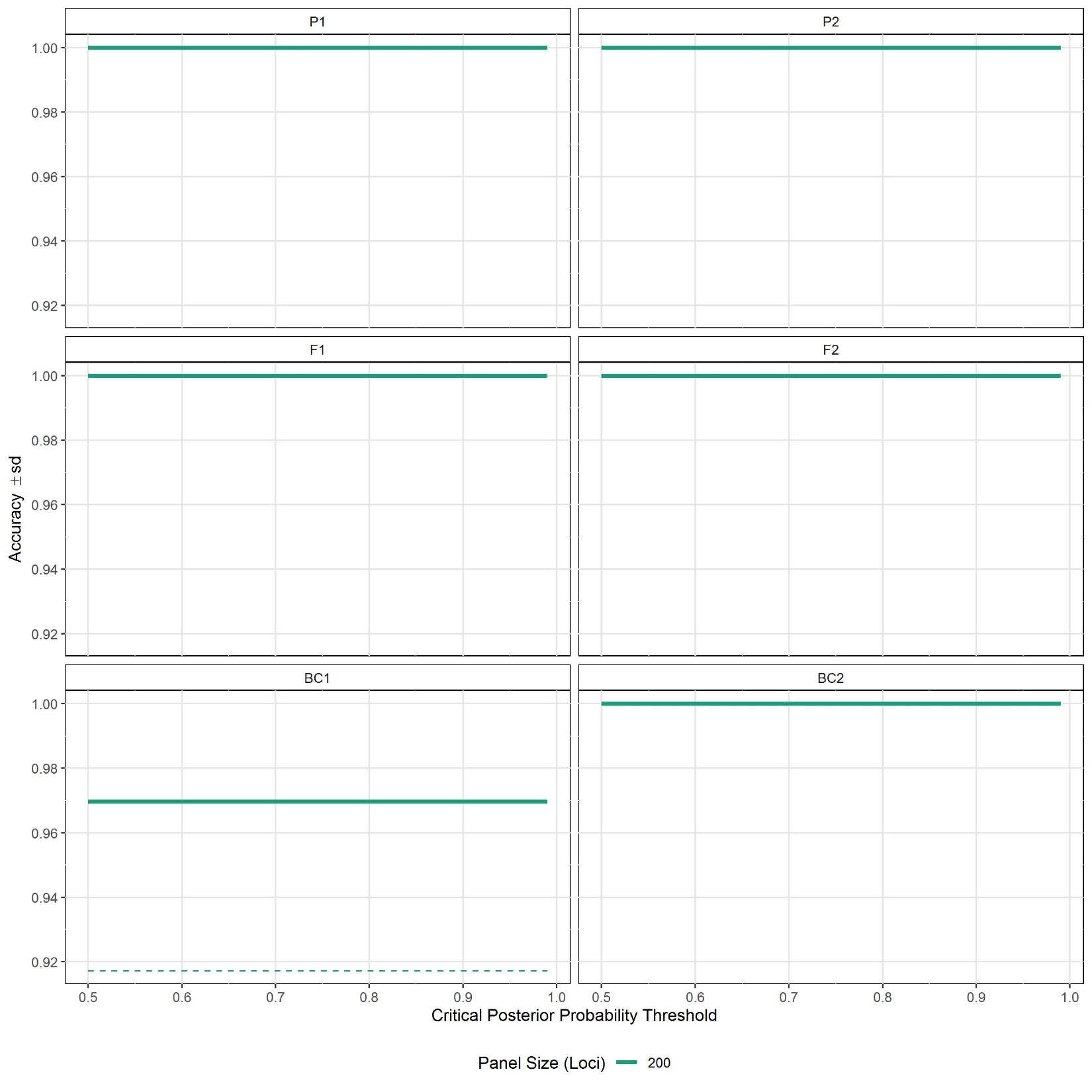

**SUPPLEMENT S14** Power analysis for detecting genotype frequency classes via HYBRIDDETECTIVE workflow. Results are shown here for *Luxilus pilsbryi* X *Luxilus zonatus*. P1 and P2 = pure; F1 and F2 = first filial and second filial hybrids; BC1 and BC2 = backcrosses.

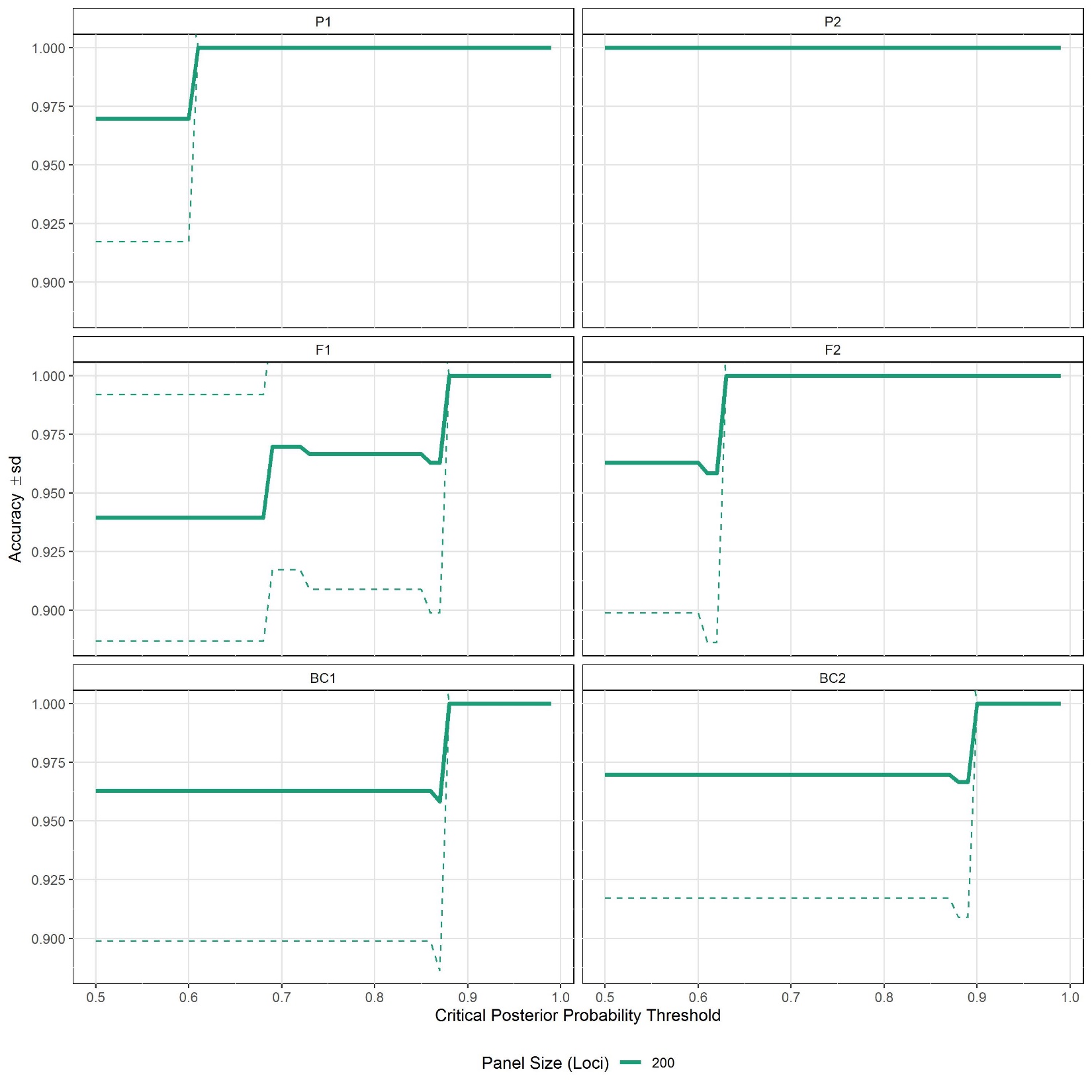

**SUPPLEMENT S15** Power analysis for detecting genotype frequency classes via HYBRIDDETECTIVE workflow. Results are shown here for *Luxilus pilsbryi* X *Lythrurus umbratilis*. P1 and P2 = pure; F1 and F2 = first filial and second filial hybrids; BC1 and BC2 = backcrosses.

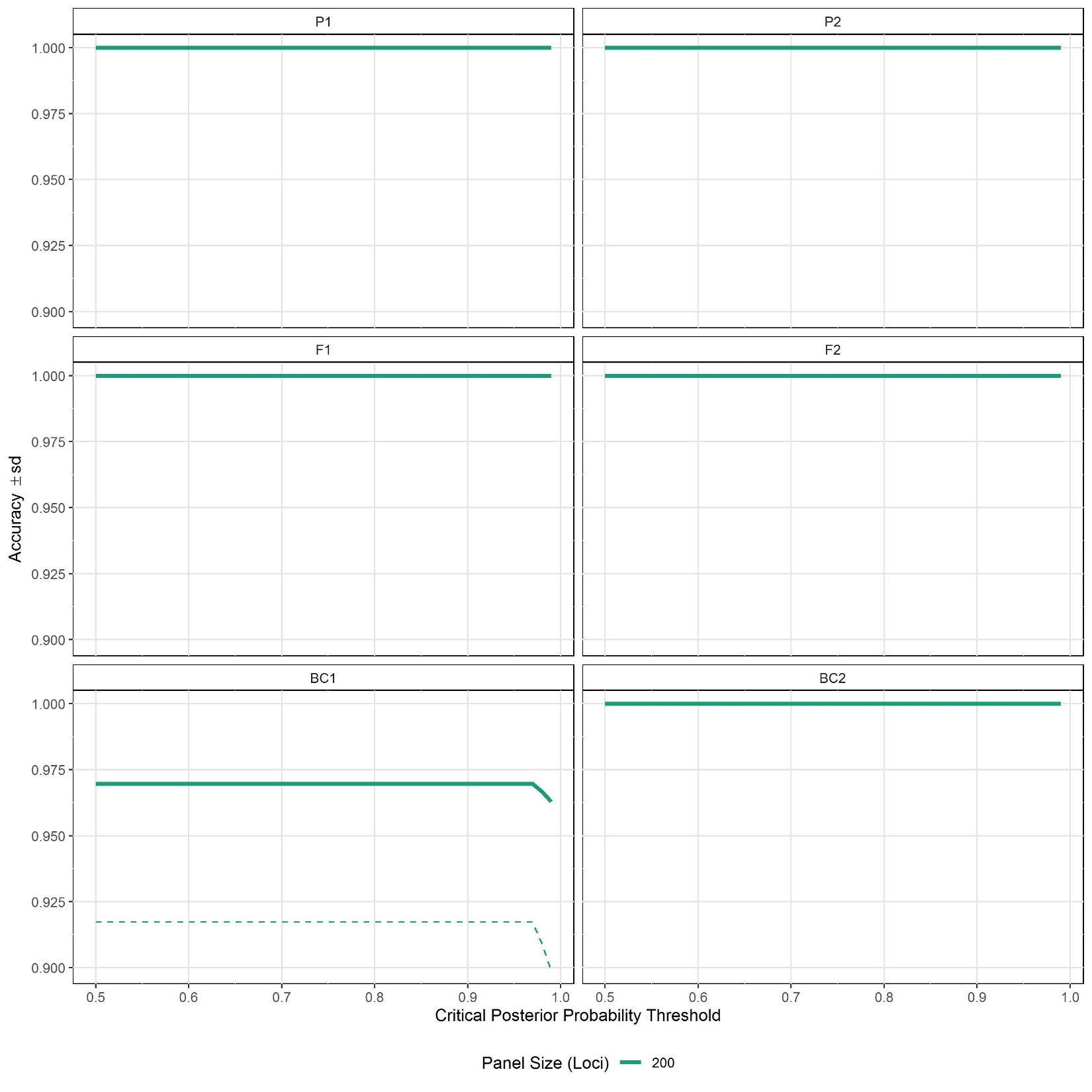

**SUPPLEMENT S16** Power analysis for detecting genotype frequency classes via HYBRIDDETECTIVE workflow. Results are shown here for *Luxilus pilsbryi* X *Notropis percobromus*. P1 and P2 = pure; F1 and F2 = first filial and second filial hybrids; BC1 and BC2 = backcrosses.

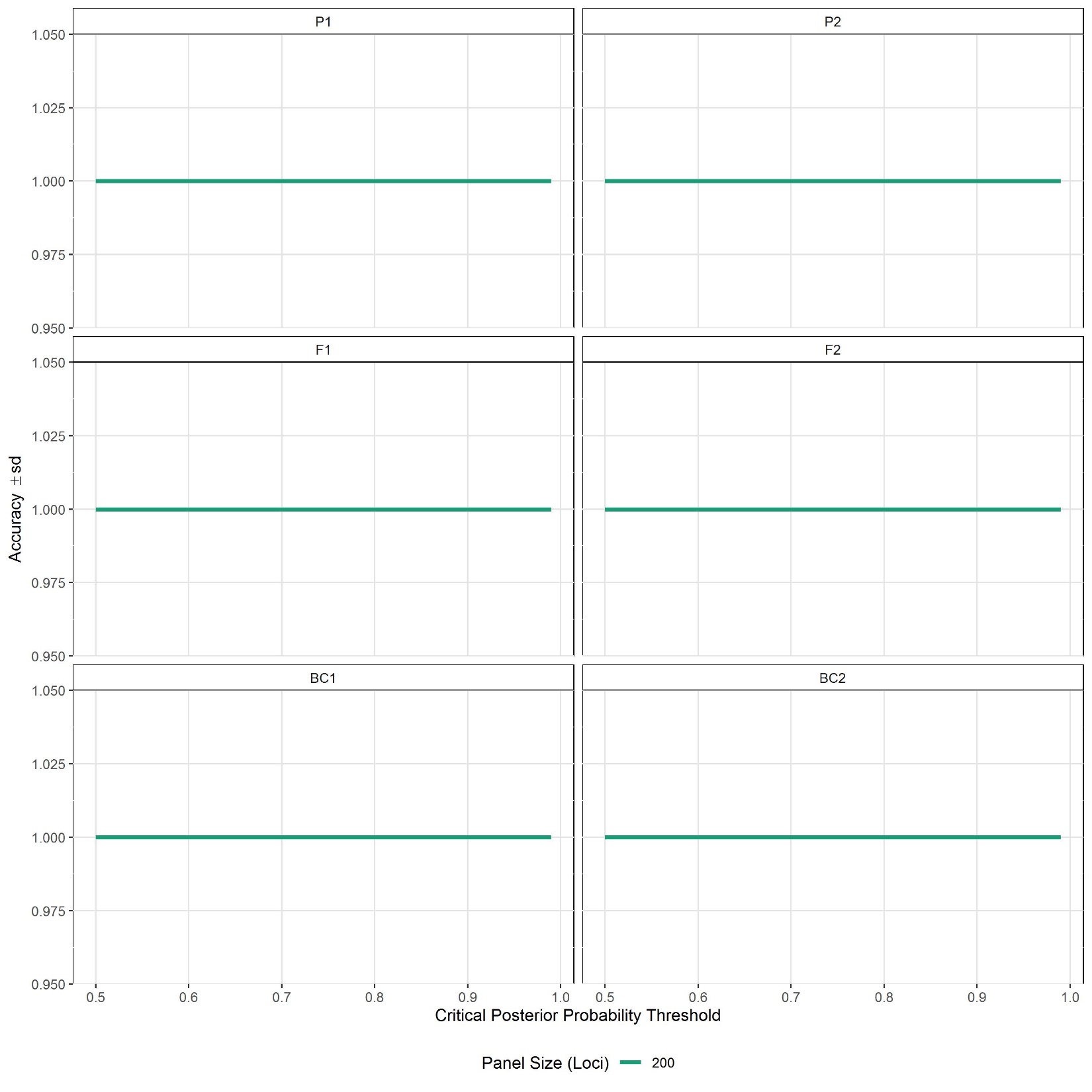

**SUPPLEMENT S17** Power analysis for detecting genotype frequency classes via HYBRIDDETECTIVE workflow. Results are shown here for *Luxilus zonatusi* X *Pimephales notatus*. P1 and P2 = pure; F1 and F2 = first filial and second filial hybrids; BC1 and BC2 = backcrosses.

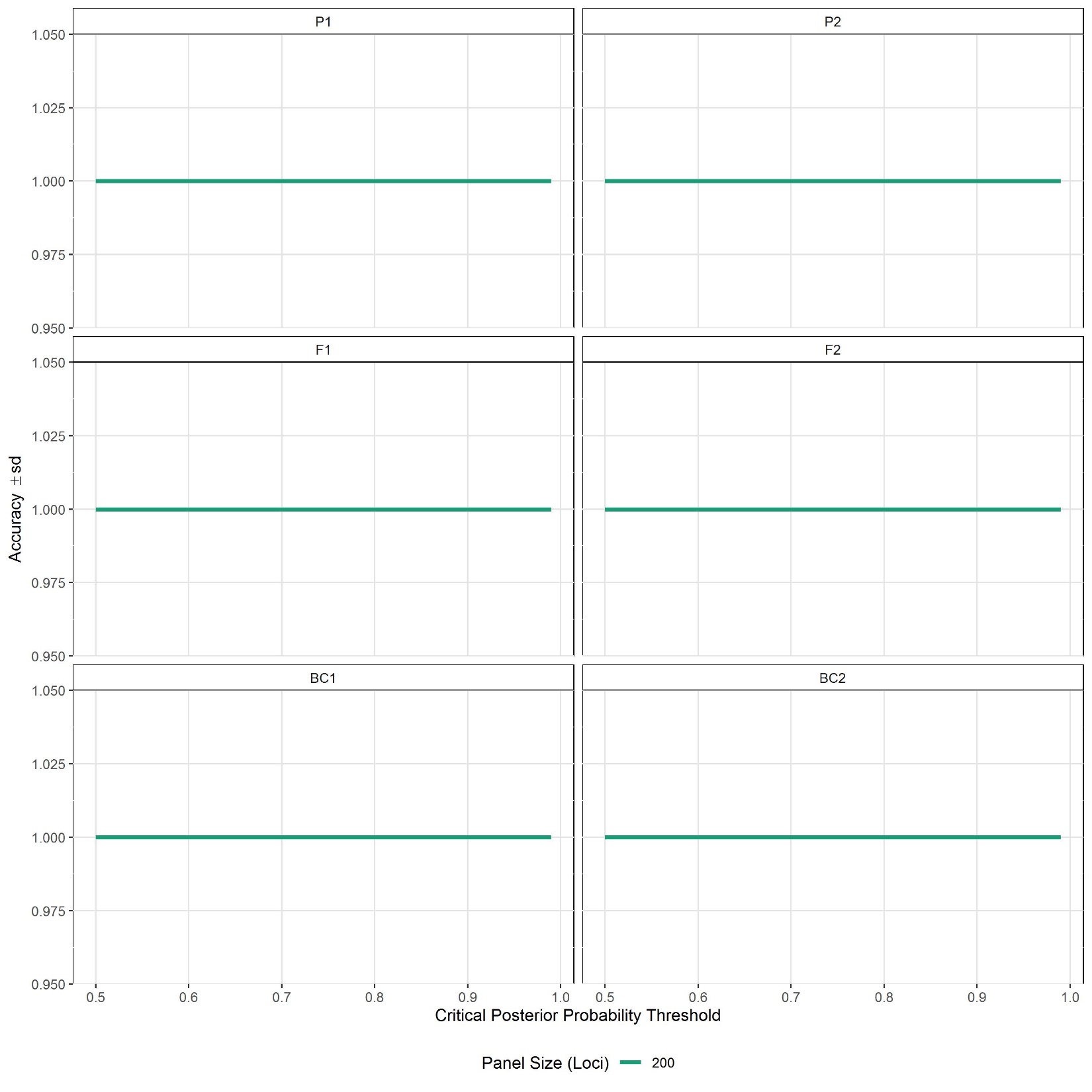

**SUPPLEMENT S18** Power analysis for detecting genotype frequency classes via HYBRIDDETECTIVE workflow. Results are shown here for *Notropis boops* X *Notropis nubilus*. P1 and P2 = pure; F1 and F2 = first filial and second filial hybrids; BC1 and BC2 = backcrosses.

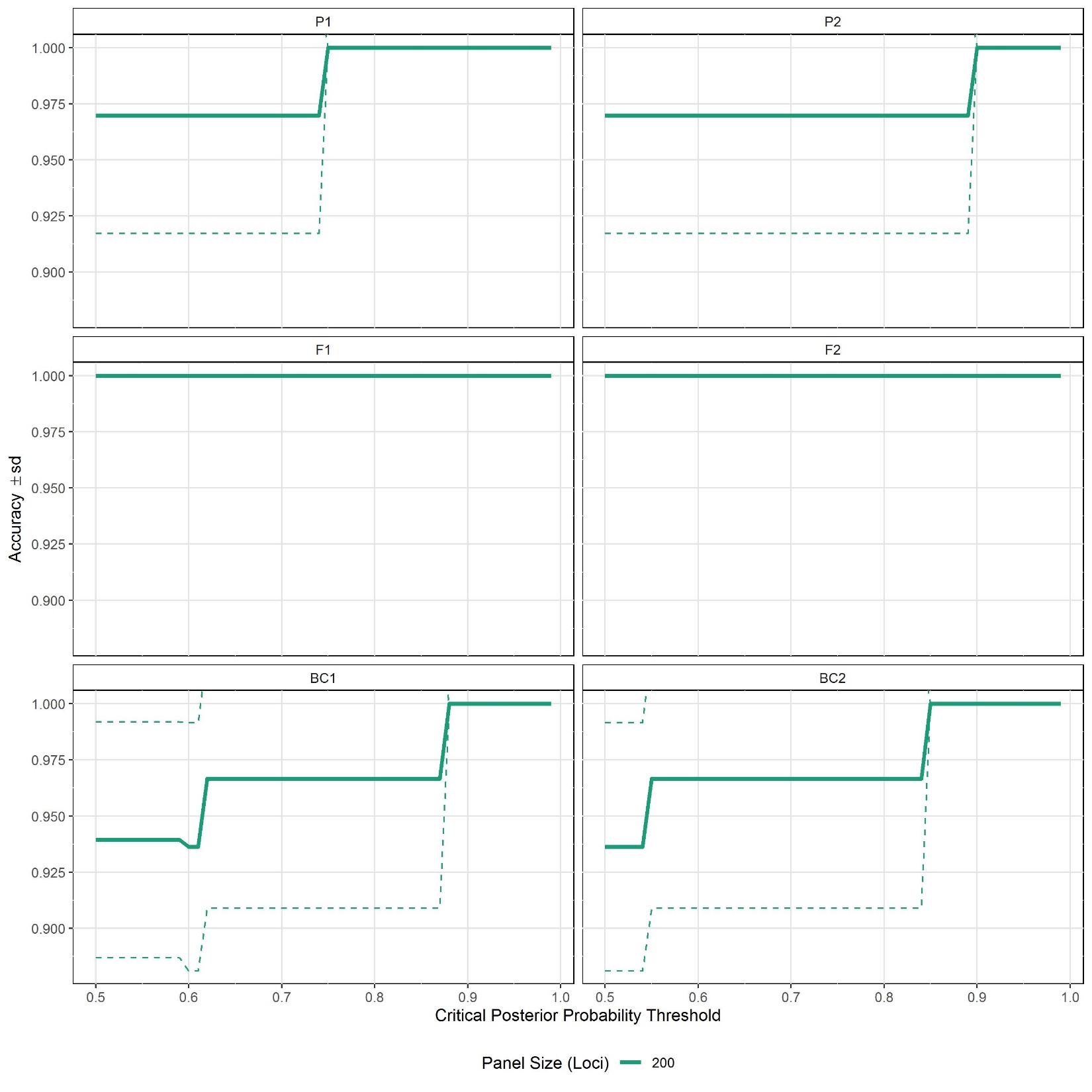

**SUPPLEMENT S19** Power analysis for detecting genotype frequency classes via HYBRIDDETECTIVE workflow. Results are shown here for *Pimephales notatus* X *Semotilus atromaculatus*. P1 and P2 = pure; F1 and F2 = first filial and second filial hybrids; BC1 and BC2 = backcrosses.

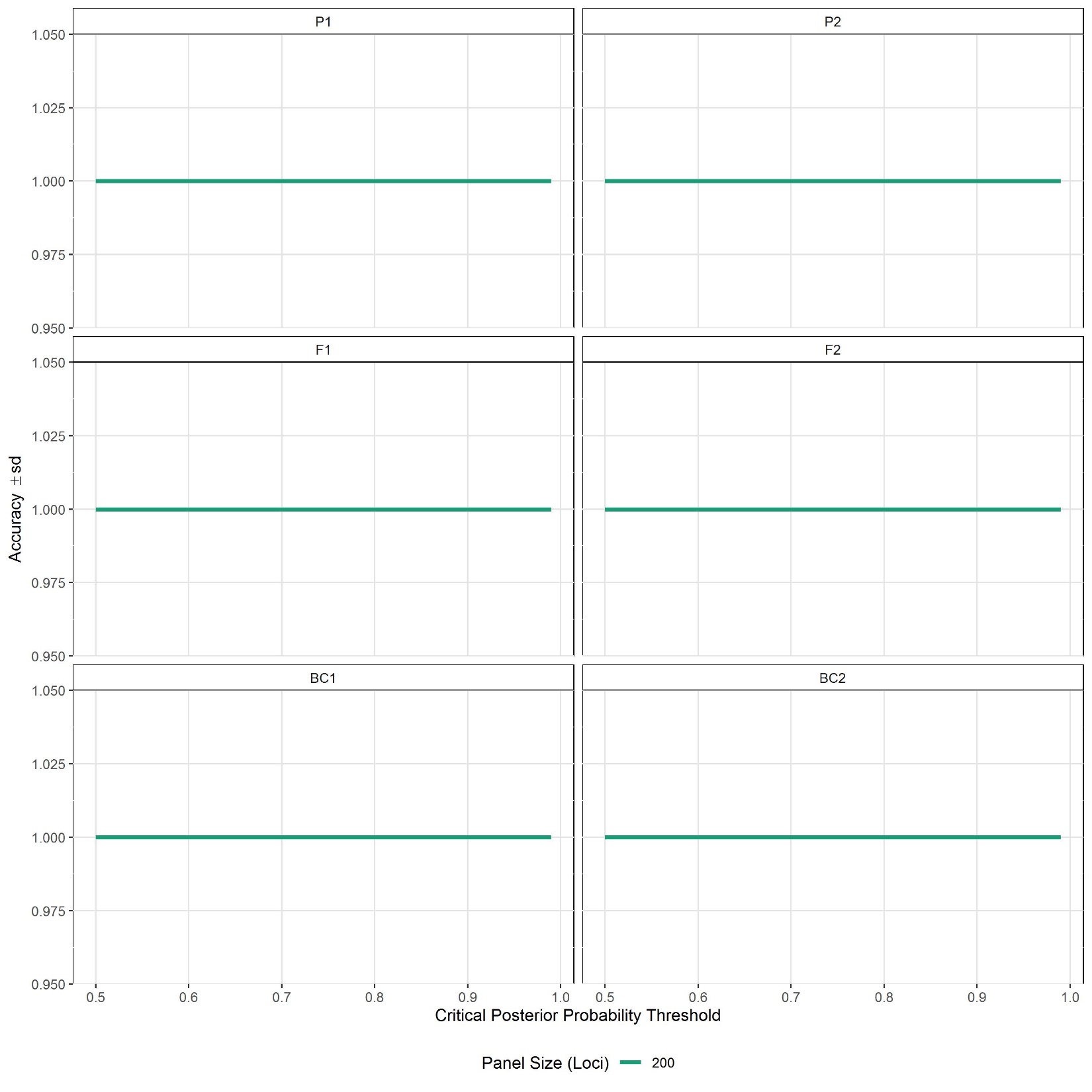

**SUPPLEMENT S20** List of 18 unique hybridizing fish species pairs observed in the White River Basin, USA. Google Scholar was used to search existing literature. Each pair was searched using "hybrid" and the two specific epithets for the pair (e.g., +hybrid* anomalum oligolepis). Searches were also conducted based on related species. Corush et al. (2021) conducted a literature review and compiled known hybrids from museum records and provided an invaluable resource here. Note this list does not include 8 of the 70 total putative hybrid individuals found: multispecies hybrids (more than two ancestral species detected).

| No. | Species A | Species B | N Indiv. | Literature Match | Species | Reference |
| --- | --- | --- | --- | --- | --- | --- |
| 1 | *Campostoma anomalum* | *Campostoma oligolepis* | 29 | yes | *C. anomalum* x *C. oligolepis* | Rakocinski, 1980 |
| 2 | *Campostoma anomalum* | *Chrosomus erythrogaster* | 1 | yes | *C. anomalum* x *C. erythrogaster* | Grady & Cashner, 1988 |
| 3 | *Campostoma anomalum* | *Luxilus pilsbryi* | 1 | related | *C. anomalum* x *L. chrysocephalus* | Grady & Cashner, 1988 |
| 4 | *Campostoma oligolepis* | *Notropis telescopus* | 1 | no | - | - |
| 5 | *Cyprinella galactura* | *Cyprinella whipplei* | 2 | yes | *C. galactura* x *C. whipplei* | Corush et al., 2021 |
| 6 | *Cyprinella whipplei* | *Lythrurus umbratilis* | 1 | no | - | - |
| 7 | *Luxilus chrysocephalus* | *Luxilus zonatus* | 2 | yes | *L. chrysocepalus* x *L. zonatus* | Corush et al., 2021 |
| 8 | *Luxilus chrysocephalus* | *Semotilus atromaculatus* | 1 | yes | *L. chrysocepalus* x *S. atromaculatus* | Corush et al., 2021 |
| 9 | *Luxilus pilsbryi* | *Lythrurus umbratilis* | 6 | related | *L. chrysocephlus* x *L. umbratilis* | Corush et al., 2021 |
| 10 | *Luxilus pilsbryi* | *Notropis percobromus* | 2 | yes | *L. pilsbryi* x *N. rubellus* | Cross 1954 |
| 11 | *Luxilus pilsbryi* | *Luxilus zonatus* | 8 | related | *L. zonatus* x *L. chrysocephalus* | Corush et al., 2021 |
| 12 | *Luxilus pilsbryi* | *Luxilus chrysocephalus* | 1 | yes | *L. pilsbryi* x *L. chrysocepalus* | Corush et al., 2021 |
| 13 | *Luxilus zonatus* | *Pimephales notatus* | 1 | no | - | - |
| 14 | *Notropis boops* | *Notropis nubilus* | 2 | yes | *N. boops* x *N. nubilus* | Corush et al., 2021 |
| 15 | *Pimephales notatus* | *Semotilus atromaculatus* | 1 | no | - | - |
| 16 | *Micropterus salmoides* | *Micropterus dolomieu* | 1 | yes | *M. salmoides* x *M. dolomieu* | Barthel et al., 2010 |
| 17 | *Etheostoma spectabile* | *Etheostoma caeruleum* | 1 | yes | *E. spectabile* x *E. caeruleum* | Keck & Near, 2009 |
| 18 | *Etheostoma juliae* | *Etheostoma zonale* | 1 | no | - | - |

**SUPPLEMENT S21** Summary of *N*=70 hybrid individuals form the White River Basin of the Ozarks. Species codes provided below.

| INDIVIDUAL | SPECIES | LATITUDE | LONGITUDE | INFERRED PARENTAL PAIR | sNMF | SNAPCLUST | DAPC | NEWHYBRIDS | POSTERIOR |
| --- | --- | --- | --- | --- | --- | --- | --- | --- | --- |
| Z01CAMANO01 | *Campostoma anomalum* | 35.90463 | -91.63537 | CAMANO-CAMOLI | CAMANO-CAMOLI | PURE | PURE | PURE | 100% |
| Z02CAMANO01 | *Campostoma anomalum* | 36.09227 | -91.75397 | CAMANO-CAMOLI | CAMANO-CAMOLI | PURE | PURE | BC1 | 89% |
| Z03CAMOLI01 | *Campostoma oligolepis* | 36.20048 | -91.759 | CAMANO-CAMOLI | CAMANO-CAMOLI | CAMANO-CAMOLI | CAMANO | F1 | 100% |
| Z06CAMANO01 | *Campostoma anomalum* | 36.27342 | -91.33447 | CAMANO-CAMOLI | CAMANO-CAMOLI | PURE | PURE | PURE | 100% |
| Z10CAMANO01 | *Campostoma anomalum* | 35.61367 | -91.60728 | CAMANO-CAMOLI | CAMANO-CAMOLI | PURE | PURE | BC1 | 72% |
| Z10CAMANO02 | *Campostoma anomalum* | 35.61367 | -91.60728 | CAMANO-CAMOLI | CAMANO-CAMOLI | CAMANO-CAMOLI | PURE | BC1 | 100% |
| Z11CAMANO01 | *Campostoma anomalum* | 35.5444 | -91.56472 | CAMANO-CAMOLI | CAMANO-CAMOLI | PURE | PURE | PURE | 100% |
| Z11CAMANO02 | *Campostoma anomalum* | 35.5444 | -91.56472 | CAMANO-CAMOLI | CAMANO-CAMOLI | PURE | PURE | PURE | 100% |
| Z11CAMANO03 | *Campostoma anomalum* | 35.5444 | -91.56472 | CAMANO-CAMOLI | CAMANO-CAMOLI | PURE | PURE | PURE | 100% |
| Z18CAMANO01 | *Campostoma anomalum* | 35.54105 | -91.77822 | CAMANO-CAMOLI | CAMANO-CAMOLI | PURE | PURE | PURE | 100% |
| Z18CAMANO02 | *Campostoma anomalum* | 35.54105 | -91.77822 | CAMANO-CAMOLI | CAMANO-CAMOLI | PURE | PURE | PURE | 100% |
| Z24CAMANO05 | *Campostoma anomalum* | 36.14685 | -92.07063 | CAMANO-CAMOLI | CAMANO-CAMOLI | CAMANO-CAMOLI | CAMANO-CAMOLI | F1 | 99% |
| Z25CAMANO01 | *Campostoma anomalum* | 36.33653 | -92.47492 | CAMANO-CAMOLI | CAMANO-CAMOLI | PURE | PURE | PURE | 100% |
| Z27CAMANO01 | *Campostoma anomalum* | 36.06698 | -92.63615 | CAMANO-CAMOLI | CAMANO-CAMOLI | PURE | PURE | PURE | 100% |
| Z29CAMOLI01 | *Campostoma oligolepis* | 36.02182 | -92.67905 | CAMANO-CAMOLI | CAMANO-CAMOLI | CAMANO-CAMOLI | PURE | BC2 | 100% |
| Z33CAMANO02 | *Campostoma anomalum* | 36.23702 | -92.98725 | CAMANO-CAMOLI | CAMANO-CAMOLI | CAMANO-CAMOLI | PURE | BC1 | 100% |
| Z33CAMOLI02 | *Campostoma oligolepis* | 36.23702 | -92.98725 | CAMANO-CAMOLI | CAMANO-CAMOLI | CAMANO-CAMOLI | PURE | F1 | 100% |
| Z34CAMOLI02 | *Campostoma oligolepis* | 36.15450 | -92.90213 | CAMANO-CAMOLI | CAMANO-CAMOLI | CAMANO-CAMOLI | CAMANO | F1 | 100% |
| Z36CAMANO01 | *Campostoma anomalum* | 35.94370 | -93.06555 | CAMANO-CAMOLI | CAMANO-CAMOLI | PURE | PURE | PURE | 100% |
| Z36CAMOLI01 | *Campostoma oligolepis* | 35.94370 | -93.06555 | CAMANO-CAMOLI | CAMANO-CAMOLI | CAMANO-CAMOLI | PURE | F1 | 100% |
| Z37CAMOLI04 | *Campostoma oligolepis* | 35.96167 | -93.23573 | CAMANO-CAMOLI | CAMANO-CAMOLI | CAMANO-CAMOLI | PURE | BC2 | 100% |
| Z37CAMOLI05 | *Campostoma oligolepis* | 35.96167 | -93.23573 | CAMANO-CAMOLI | CAMANO-CAMOLI | CAMANO-CAMOLI | PURE | BC2 | 95% |
| Z42CAMANO09 | *Campostoma anomalum* | 36.13102 | -93.94767 | CAMANO-CAMOLI | CAMANO-CAMOLI | CAMANO-CAMOLI | PURE | BC1 | 100% |
| Z47CAMANO01 | *Campostoma anomalum* | 36.42222 | -93.40465 | CAMANO-CAMOLI | CAMANO-CAMOLI | CAMANO-CAMOLI | PURE | BC1 | 100% |
| Z58CAMANO01 | *Campostoma anomalum* | 37.45613 | -91.69537 | CAMANO-CAMOLI | CAMANO-CAMOLI | PURE | PURE | PURE | 100% |
| Z67CAMANO01 | *Campostoma anomalum* | 36.88635 | -92.47363 | CAMANO-CAMOLI | CAMANO-CAMOLI | CAMANO-CAMOLI | PURE | BC1 | 88% |
| Z68CAMANO01 | *Campostoma anomalum* | 37.02342 | -92.76715 | CAMANO-CAMOLI | CAMANO-CAMOLI | PURE | PURE | PURE | 100% |
| Z71CAMANO01 | *Campostoma anomalum* | 37.18145 | -93.37052 | CAMANO-CAMOLI | CAMANO-CAMOLI | PURE | PURE | PURE | 100% |
| Z55CAMANO06 | *Campostoma anomalum* | 37.35483 | -90.97105 | CAMANO-CHRERY | CAMANO-CHRERY | CAMANO-CHRERY | PURE | F1 | 100% |
| Z52CAMANO02 | *Campostoma anomalum* | 36.82858 | -90.81385 | CAMANO-LUXPIL | CAMANO-LUXPIL | CAMANO-CHRERY | PURE | F2 | 100% |
| Z69CAMOLI04 | *Campostoma oligolepis* | 37.04205 | -93.13870 | CAMOLI-LUXPIL-NOTNUB | CAMOLI-LUXPIL-NOTNUB | LYTUMB-CAMOLI | CAMOLI-NOTNUB | NA | NA |
| Z04CAMOLI04 | *Campostoma oligolepis* | 36.38583 | -91.80977 | CAMOLI-NOTTEL | CAMOLI-NOTTEL | LYTUMB-CAMOLI | PURE | F1 | 96% |
| Z06CYPGAL02 | *Cyprinella galactura* | 36.27342 | -91.33447 | CYPGAL-CYPWHI | CYPGAL-CYPWHI | CYPGAL-CYPWHI | PURE | BC1 | 91% |
| Z06CYPGAL07 | *Cyprinella galactura* | 36.27342 | -91.33447 | CYPGAL-CYPWHI | CYPGAL-CYPWHI | CYPGAL-CYPWHI | PURE | F1 | 99% |
| Z41CYPWHI02 | *Cyprinella galactura* | 36.04162 | -93.70483 | CYPWHI-LYTUMB | CYPWHI-LYTUMB | LYTUMB-CYPWHI | PURE | BC2 | 100% |
| Z48ETHJUL05 | *Etheostoma juliae* | 36.25182 | -93.44542 | ETHJUL-ETHZON | ETHJUL-ETHZON | ETHJUL-ETHFLA | PURE | BC2 | 100% |
| Z01ETHSPE20 | *Etheostoma spectabile* | 35.90463 | -91.63537 | ETHSPE-ETHCAE | ETHSPE-ETHCAE | ETHSPE-ETHUNI | PURE | BC2 | 100% |
| Z41LUXCHR10 | *Luxilus chrysocephalus* | 36.04162 | -93.70483 | LUXCHR-LUXPIL-CYPGAL | LUXCHR-LUXPIL-CYPGAL | LYTUMB-LUXCHR | PURE | NA | NA |
| Z52LUXCHR03 | *Luxilus chrysocephalus* | 36.82858 | -90.81385 | LUXCHR-LUXZON | LUXCHR-LUXZON | LYTUMB-LUXCHR | LUXZON | F1 | 100% |
| Z62LUXCHR02 | *Luxilus chrysocephalus* | 36.55238 | -91.53942 | LUXCHR-LUXZON | LUXCHR-LUXZON | PURE | PURE | PURE | 100% |
| Z05LUXCHR01 | *Luxilus chrysocephalus* | 36.44377 | -91.67055 | LUXCHR-NOTNUB-LUXPIL | LUXCHR-NOTNUB-LUXPIL | LYTUMB-LUXCHR | PURE | NA | NA |
| Z38LUXPIL08 | *Luxilus pilsbryi* | 36.03938 | -93.33633 | LUXPIL-LUXCHR | LUXPIL-LUXCHR | LUXPIL-LYTUMB | PURE | BC1 | 99% |
| Z31LUXPIL02 | *Luxilus pilsbryi* | 35.92747 | -92.89285 | LUXPIL-LUXZON | LUXPIL-LUXZON | PURE | PURE | PURE | 100% |
| Z32LUXPIL02 | *Luxilus pilsbryi* | 36.45005 | -93.07547 | LUXPIL-LUXZON | LUXPIL-LUXZON | PURE | PURE | PURE | 100% |
| Z40LUXPIL08 | *Luxilus pilsbryi* | 36.14380 | -93.59412 | LUXPIL-LUXZON | LUXPIL-LUXZON | PURE | PURE | PURE | 100% |
| Z42LUXPIL10 | *Luxilus pilsbryi* | 36.13102 | -93.94767 | LUXPIL-LUXZON | LUXPIL-LUXZON | PURE | PURE | PURE | 100% |
| Z44LUXPIL07 | *Luxilus pilsbryi* | 35.82850 | -93.83217 | LUXPIL-LUXZON | LUXPIL-LUXZON | PURE | PURE | PURE | 100% |
| Z46LUXPIL02 | *Luxilus pilsbryi* | 35.98301 | -94.17291 | LUXPIL-LUXZON | LUXPIL-LUXZON | PURE | PURE | PURE | 100% |
| Z46LUXPIL03 | *Luxilus pilsbryi* | 35.98301 | -94.17291 | LUXPIL-LUXZON | LUXPIL-LUXZON | PURE | PURE | PURE | 100% |
| Z51LUXZON06 | *Luxilus zonatus* | 36.57930 | -91.04640 | LUXPIL-LUXZON | LUXPIL-LUXZON | PURE | PURE | PURE | 100% |
| Z24LUXPIL06 | *Luxilus pilsbryi* | 36.14685 | -92.07063 | LUXPIL-NOTPER | LUXPIL-NOTPER | LUXPIL-LYTUMB | PURE | BC1 | 99% |
| Z31LUXPIL03 | *Luxilus pilsbryi* | 35.92747 | -92.89285 | LUXPIL-NOTPER | LUXPIL-NOTPER | LUXPIL-LYTUMB | PURE | F1 | 100% |
| Z58LUXZON06 | *Luxilus zonatus* | 37.45613 | -91.69537 | LUXZON-PIMNOT | LUXZON-PIMNOT | LYTUMB-LUXZON | PURE | F1 | 100% |
| Z52LYTUMB10 | *Lythrurus umbratilis* | 36.82858 | -90.81385 | LYTUMB-NOTNUB-LUXPIL | LYTUMB-NOTNUB-LUXPIL | LYTUMB-LUXCHR | PURE | NA | NA |
| Z15MICSAL03 | *Micropterus salmoides* | 34.36425 | -91.23615 | MICSAL-MICDOL | MICSAL-MICDOL | MICSAL-MICDOL | PURE | F1 | 100% |
| Z19CAMOLI02 | *Campostoma oligolepis* | 35.73682 | -92.10778 | CAMANO-CAMOLI | PURE | CAMANO-CAMOLI | PURE | BC2 | 99% |
| Z24LUXPIL01 | *Luxilus pilsbryi* | 36.14685 | -92.07063 | LUXPIL-LYTUMB | PURE | LUXPIL-LYTUMB | PURE | BC1 | 98% |
| Z29LUXPIL06 | *Luxilus pilsbryi* | 36.02182 | -92.67905 | LUXPIL-LYTUMB | PURE | LUXPIL-LYTUMB | NOTNUB | BC1 | 95% |
| Z35LUXPIL05 | *Luxilus pilsbryi* | 35.98273 | -93.04035 | LUXPIL-LYTUMB | PURE | LUXPIL-LYTUMB | PURE | BC1 | 100% |
| Z46LUXPIL10 | *Luxilus pilsbryi* | 35.98301 | -94.17291 | LUXPIL-LYTUMB | PURE | LUXPIL-LYTUMB | NOTNUB | F2 | 99% |
| Z48LUXPIL04 | *Luxilus pilsbryi* | 36.25182 | -93.44542 | LUXPIL-LYTUMB | PURE | LUXPIL-LYTUMB | PURE | BC1 | 92% |
| Z58LUXCHR02 | *Luxilus chrysocephalus* | 37.45613 | -91.69537 | LUXCHR-SEMATR | PURE | LUXCHR-SEMATR | PURE | BC1 | 100% |
| Z67LUXPIL02 | *Luxilus pilsbryi* | 36.88635 | -92.47363 | LUXPIL-LYTUMB | PURE | LUXPIL-LYTUMB | NOTNUB | BC1 | 86% |
| Z48NOTBOO03 | *Notropis boops* | 36.25182 | -93.44542 | NOTBOO-LUXPIL-PIMNOT | NOTBOO-LUXPIL-PIMNOT | LYTUMB-NOTBOO | PURE | NA | NA |
| Z07NOTBOO09 | *Notropis boops* | 35.8711 | -91.31063 | NOTBOO-NOTNUB | NOTBOO-NOTNUB | PURE | PURE | PURE | 100% |
| Z41NOTBOO06 | *Notropis boops* | 36.04162 | -93.70483 | NOTBOO-NOTNUB | NOTBOO-NOTNUB | NOTBOO-NOTNUB | PURE | BC1 | 95% |
| Z04NOTMAY01 | *Noturus maydeni* | 36.38583 | -91.80977 | NOTMAY-NOTEXI-NOTALB | NOTMAY-NOTEXI-NOTALB | PURE | PURE | NA | NA |
| Z61NOTNUB11 | *Notropis nubilus* | 36.85519 | -91.67677 | NOTNUB-LUXPIL-NOTPER | NOTNUB-LUXPIL-NOTPER | NOTNUB-SEMATR | PURE | NA | NA |
| Z51NOTNUB06 | *Notropis nubilus* | 36.57930 | -91.04640 | NOTNUB-LUXPIL-SEMATR | NOTNUB-LUXPIL-SEMATR | NOTNUB-SEMATR | PURE | NA | NA |
| Z43PIMNOT02 | *Pimephales notatus* | 36.04882 | -93.97490 | PIMNOT-SEMATR | PIMNOT-SEMATR | PIMNOT-SEMATR | PURE | F2 | 93% |

**SUPPLEMENT S22** Intra-familial species comparisons from the study of hybridization across the White River Basin. For each species pair (N=137) the table shows the number of times they co-occurred together in N=75 community samples, the number of hybrid individuals detected, and the Weir and Cockerham's *F*_ST_ among them based on SNP genotypes. Pairs are grouped by families. Species codes = First three letters of Genus + first three of specific epithet (full list below).

| **Family** | **Sp1** | **Sp2** | **Cooccurrences** | **Hybrids** | **FstWC** |  | **Family** | **Sp1** | **Sp2** | **Cooccurrences** | **Hybrids** | **FstWC** |
| --- | --- | --- | --- | --- | --- | --- | --- | --- | --- | --- | --- | --- |
| Centrarchidae | MICDOL | MICSAL | 5 | 1 | 0.8464 |  | Leuciscidae | LUXCHR | NOTBOO | 7 | 0 | 0.8622 |
| Centrarchidae | LEPMAC | MICDOL | 7 | 0 | 0.9769 |  | Leuciscidae | CYPGAL | NOTPER | 7 | 0 | 0.928 |
| Centrarchidae | LEPMEG | MICSAL | 10 | 0 | 0.9618 |  | Leuciscidae | LUXCHR | NOTTEL | 7 | 0 | 0.9108 |
| Centrarchidae | LEPMAC | MICSAL | 12 | 0 | 0.9737 |  | Leuciscidae | LUXZON | NOTTEL | 7 | 0 | 0.9132 |
| Centrarchidae | LEPMAC | LEPMEG | 17 | 0 | 0.9081 |  | Leuciscidae | NOTPER | NOTTEL | 7 | 0 | 0.929 |
| Centrarchidae | LEPMEG | MICDOL | 25 | 0 | 0.9639 |  | Leuciscidae | CYPGAL | PIMNOT | 7 | 0 | 0.9361 |
| Cottidae | COTCAR | COTHYP | 3 | 0 | 0.8166 |  | Leuciscidae | PIMNOT | SEMATR | 7 | 1 | 0.9717 |
| Fundulidae | FUNCAT | FUNOLI | 16 | 0 | 0.9773 |  | Leuciscidae | CAMOLI | CYPWHI | 8 | 0 | 0.9537 |
| Ictaluridae | NOTALB | NOTMAY | 0 | 0 | 0.659 |  | Leuciscidae | LUXZON | NOTNUB | 8 | 0 | 0.8643 |
| Ictaluridae | NOTALB | NOTEXI | 1 | 0 | 0.9082 |  | Leuciscidae | LUXCHR | PIMNOT | 8 | 0 | 0.924 |
| Ictaluridae | NOTEXI | NOTMAY | 1 | 0 | 0.8699 |  | Leuciscidae | LUXPIL | PIMNOT | 8 | 0 | 0.8919 |
| Leuciscidae | CHRERY | CYPWHI | 0 | 0 | 0.9846 |  | Leuciscidae | CYPGAL | LUXPIL | 9 | 0 | 0.887 |
| Leuciscidae | LUXPIL | LUXZON | 0 | 8 | 0.6467 |  | Leuciscidae | LUXCHR | LUXPIL | 9 | 1 | 0.7996 |
| Leuciscidae | CYPWHI | LYTUMB | 0 | 1 | 0.9518 |  | Leuciscidae | CYPGAL | NOTNUB | 9 | 0 | 0.8954 |
| Leuciscidae | CHRERY | NOTPER | 0 | 0 | 0.9745 |  | Leuciscidae | NOTNUB | NOTPER | 9 | 0 | 0.8814 |
| Leuciscidae | CYPWHI | NOTPER | 0 | 0 | 0.9373 |  | Leuciscidae | LUXPIL | NOTTEL | 9 | 0 | 0.8706 |
| Leuciscidae | LYTUMB | NOTPER | 0 | 0 | 0.921 |  | Leuciscidae | CAMANO | CHRERY | 10 | 1 | 0.9624 |
| Leuciscidae | LYTUMB | SEMATR | 0 | 0 | 0.9905 |  | Leuciscidae | CAMANO | CYPGAL | 10 | 0 | 0.9636 |
| Leuciscidae | CHRERY | CYPGAL | 1 | 0 | 0.9834 |  | Leuciscidae | CAMOLI | CYPGAL | 10 | 0 | 0.9649 |
| Leuciscidae | CYPGAL | CYPWHI | 1 | 2 | 0.7973 |  | Leuciscidae | CAMOLI | LUXZON | 10 | 0 | 0.9521 |
| Leuciscidae | CHRERY | LUXCHR | 1 | 0 | 0.9637 |  | Leuciscidae | LUXPIL | NOTBOO | 10 | 0 | 0.8678 |
| Leuciscidae | CYPWHI | LUXCHR | 1 | 0 | 0.9167 |  | Leuciscidae | CAMANO | NOTPER | 10 | 0 | 0.9574 |
| Leuciscidae | CYPWHI | LUXZON | 1 | 0 | 0.9266 |  | Leuciscidae | LUXPIL | NOTPER | 10 | 2 | 0.8599 |
| Leuciscidae | CAMOLI | LYTUMB | 1 | 0 | 0.951 |  | Leuciscidae | CAMOLI | NOTTEL | 10 | 1 | 0.9693 |
| Leuciscidae | CYPGAL | LYTUMB | 1 | 0 | 0.9375 |  | Leuciscidae | LUXPIL | SEMATR | 10 | 0 | 0.9094 |
| Leuciscidae | LUXCHR | LYTUMB | 1 | 0 | 0.8745 |  | Leuciscidae | CYPGAL | NOTBOO | 11 | 0 | 0.9104 |
| Leuciscidae | LUXPIL | LYTUMB | 1 | 6 | 0.8118 |  | Leuciscidae | CAMOLI | NOTPER | 11 | 0 | 0.9642 |
| Leuciscidae | CHRERY | NOTTEL | 1 | 0 | 0.9752 |  | Leuciscidae | CAMANO | SEMATR | 11 | 0 | 0.9375 |
| Leuciscidae | CYPWHI | NOTTEL | 1 | 0 | 0.9486 |  | Leuciscidae | CAMANO | LUXZON | 12 | 0 | 0.9487 |
| Leuciscidae | LYTUMB | NOTTEL | 1 | 0 | 0.9354 |  | Leuciscidae | LUXCHR | NOTNUB | 12 | 0 | 0.8447 |
| Leuciscidae | CYPWHI | SEMATR | 1 | 0 | 0.9792 |  | Leuciscidae | CAMOLI | SEMATR | 12 | 0 | 0.9528 |
| Leuciscidae | CYPGAL | LUXZON | 2 | 0 | 0.9282 |  | Leuciscidae | NOTNUB | SEMATR | 12 | 0 | 0.9312 |
| Leuciscidae | CHRERY | LYTUMB | 2 | 0 | 0.9888 |  | Leuciscidae | CAMANO | NOTTEL | 13 | 0 | 0.9671 |
| Leuciscidae | LUXZON | LYTUMB | 2 | 0 | 0.8685 |  | Leuciscidae | NOTNUB | PIMNOT | 13 | 0 | 0.8978 |
| Leuciscidae | CHRERY | NOTBOO | 2 | 0 | 0.9551 |  | Leuciscidae | CAMANO | LUXCHR | 14 | 0 | 0.9455 |
| Leuciscidae | CHRERY | PIMNOT | 2 | 0 | 0.9776 |  | Leuciscidae | NOTBOO | NOTNUB | 14 | 2 | 0.7629 |
| Leuciscidae | CYPGAL | SEMATR | 2 | 0 | 0.967 |  | Leuciscidae | CAMANO | PIMNOT | 14 | 0 | 0.959 |
| Leuciscidae | NOTTEL | SEMATR | 2 | 0 | 0.9725 |  | Leuciscidae | CAMOLI | LUXCHR | 15 | 0 | 0.9495 |
| Leuciscidae | CYPWHI | LUXPIL | 3 | 0 | 0.8835 |  | Leuciscidae | CAMOLI | PIMNOT | 15 | 0 | 0.9566 |
| Leuciscidae | LYTUMB | NOTNUB | 3 | 0 | 0.8448 |  | Leuciscidae | NOTBOO | PIMNOT | 18 | 0 | 0.9035 |
| Leuciscidae | LUXZON | NOTPER | 3 | 0 | 0.9057 |  | Leuciscidae | CAMANO | NOTBOO | 19 | 0 | 0.9434 |
| Leuciscidae | LYTUMB | PIMNOT | 3 | 0 | 0.9473 |  | Leuciscidae | CAMOLI | NOTBOO | 19 | 0 | 0.9434 |
| Leuciscidae | NOTPER | PIMNOT | 3 | 0 | 0.9471 |  | Leuciscidae | CAMANO | NOTNUB | 21 | 0 | 0.9449 |
| Leuciscidae | LUXZON | SEMATR | 3 | 0 | 0.94 |  | Leuciscidae | CAMANO | LUXPIL | 22 | 1 | 0.9182 |
| Leuciscidae | NOTBOO | SEMATR | 3 | 0 | 0.939 |  | Leuciscidae | LUXPIL | NOTNUB | 23 | 0 | 0.8414 |
| Leuciscidae | CAMOLI | CHRERY | 4 | 0 | 0.9696 |  | Leuciscidae | CAMOLI | LUXPIL | 24 | 0 | 0.9327 |
| Leuciscidae | CHRERY | LUXPIL | 4 | 0 | 0.9327 |  | Leuciscidae | CAMOLI | NOTNUB | 24 | 0 | 0.942 |
| Leuciscidae | CHRERY | LUXZON | 4 | 0 | 0.9541 |  | Leuciscidae | CAMANO | CAMOLI | 29 | 29 | 0.7685 |
| Leuciscidae | CAMANO | LYTUMB | 4 | 0 | 0.9527 |  | Percidae | ETHFLA | ETHJUL | 0 | 0 | 0.9675 |
| Leuciscidae | LYTUMB | NOTBOO | 4 | 0 | 0.8643 |  | Percidae | ETHJUL | ETHSPE | 0 | 0 | 0.9391 |
| Leuciscidae | CYPWHI | NOTNUB | 4 | 0 | 0.9046 |  | Percidae | ETHJUL | ETHUNI | 0 | 0 | 0.957 |
| Leuciscidae | NOTTEL | PIMNOT | 4 | 0 | 0.9464 |  | Percidae | ETHSPE | ETHUNI | 0 | 0 | 0.7806 |
| Leuciscidae | CHRERY | SEMATR | 4 | 0 | 0.9993 |  | Percidae | ETHSPE | ETHZON | 0 | 0 | 0.9381 |
| Leuciscidae | CAMANO | CYPWHI | 5 | 0 | 0.9607 |  | Percidae | ETHBLE | ETHFLA | 1 | 0 | 0.9809 |
| Leuciscidae | CHRERY | NOTNUB | 5 | 0 | 0.945 |  | Percidae | ETHBLE | ETHSPE | 1 | 0 | 0.9557 |
| Leuciscidae | CYPWHI | PIMNOT | 5 | 0 | 0.9396 |  | Percidae | ETHBLE | ETHUNI | 2 | 0 | 0.9756 |
| Leuciscidae | NOTPER | SEMATR | 5 | 0 | 0.964 |  | Percidae | ETHUNI | ETHZON | 2 | 0 | 0.9442 |
| Leuciscidae | CYPGAL | LUXCHR | 6 | 0 | 0.915 |  | Percidae | ETHFLA | ETHSPE | 3 | 0 | 0.9368 |
| Leuciscidae | LUXCHR | LUXZON | 6 | 2 | 0.8322 |  | Percidae | ETHFLA | ETHUNI | 3 | 0 | 0.9739 |
| Leuciscidae | LUXZON | NOTBOO | 6 | 0 | 0.8824 |  | Percidae | ETHFLA | ETHZON | 3 | 0 | 0.9452 |
| Leuciscidae | LUXCHR | NOTPER | 6 | 0 | 0.9029 |  | Percidae | ETHCAE | ETHSPE | 7 | 1 | 0.8453 |
| Leuciscidae | NOTBOO | NOTPER | 6 | 0 | 0.8911 |  | Percidae | ETHCAE | ETHUNI | 7 | 0 | 0.8437 |
| Leuciscidae | CYPGAL | NOTTEL | 6 | 0 | 0.9373 |  | Percidae | ETHJUL | ETHZON | 9 | 1 | 0.9302 |
| Leuciscidae | NOTBOO | NOTTEL | 6 | 0 | 0.8986 |  | Percidae | ETHCAE | ETHFLA | 10 | 0 | 0.9135 |
| Leuciscidae | NOTNUB | NOTTEL | 6 | 0 | 0.8824 |  | Percidae | ETHBLE | ETHJUL | 10 | 0 | 0.9676 |
| Leuciscidae | LUXZON | PIMNOT | 6 | 1 | 0.9235 |  | Percidae | ETHCAE | ETHJUL | 13 | 0 | 0.9193 |
| Leuciscidae | LUXCHR | SEMATR | 6 | 1 | 0.955 |  | Percidae | ETHBLE | ETHZON | 19 | 0 | 0.9373 |
| Leuciscidae | CYPWHI | NOTBOO | 7 | 0 | 0.9043 |  | Percidae | ETHCAE | ETHZON | 23 | 0 | 0.9185 |
|  |  |  |  |  |  |  | Percidae | ETHBLE | ETHCAE | 24 | 0 | 0.9277 |

**SUPPLEMENT S23** Variables with a significant (i.e., non-zero) variable importance (VIMP) for classifying locations as ones with or without hybrid individuals using random forest classification.

| **Variable** | **Units** | **Spatial Scale** | **Original name** | **VIMP x 100** | | | **p-value** |
| --- | --- | --- | --- | --- | --- | --- | --- |
|  |  |  |  | lower | mean | upper |  |
| Number of species collected | raw number | site | NA | 0.607 | 0.804 | 1.001 | <0.000001 |
| Average annual precipitation | mm | reach | pre_mm_cyr | 0.187 | 0.317 | 0.446 | 0.000001 |
| Percent protected area extent | percent extent of coverage | reach + total upstream catchment | pac_pc_use | 0.033 | 0.219 | 0.405 | 0.010374 |
| Human impact index 1993 | index | reach | hft_ix_c93 | 0.061 | 0.189 | 0.317 | 0.001953 |
| Human impact index 2009 | index | reach | hft_ix_c09 | 0.000 | 0.137 | 0.273 | 0.024691 |
| Average road density | m/km2 | reach | rdd_mk_cav | 0.011 | 0.131 | 0.250 | 0.015924 |
| Average May precipitation | mm | reach | pre_mm_c05 | -0.004 | 0.110 | 0.223 | 0.029084 |
| Average annual snow cover percent extent | percent extent of coverage | reach + total upstream catchment | snw_pc_uyr | 0.003 | 0.046 | 0.089 | 0.017232 |
| Percent coverage of temperate deciduous forest | percent extent of coverage | reach | pnv_pc_c05 | 0.013 | 0.017 | 0.022 | <0.000001 |
| Major vegetation class | majority class | reach | pnv_cl_cmj | 0.005 | 0.010 | 0.015 | 0.000040 |
| Percent coverage of evergreen/deciduous mixed forest | percent extent of coverage | reach + total upstream catchment | pnv_pc_u08 | 0.001 | 0.009 | 0.017 | 0.018537 |
| Percent coverage of evergreen/deciduous mixed forest | percent extent of coverage | reach | pnv_pc_c08 | 0.004 | 0.009 | 0.014 | 0.000203 |

**SUPPLEMENT S24** Marginal effects plots showing the relationship between the probability of a hybrid occurring at a location and certain variables that showed a significant ability to increase the prediction capacity of such incidence.

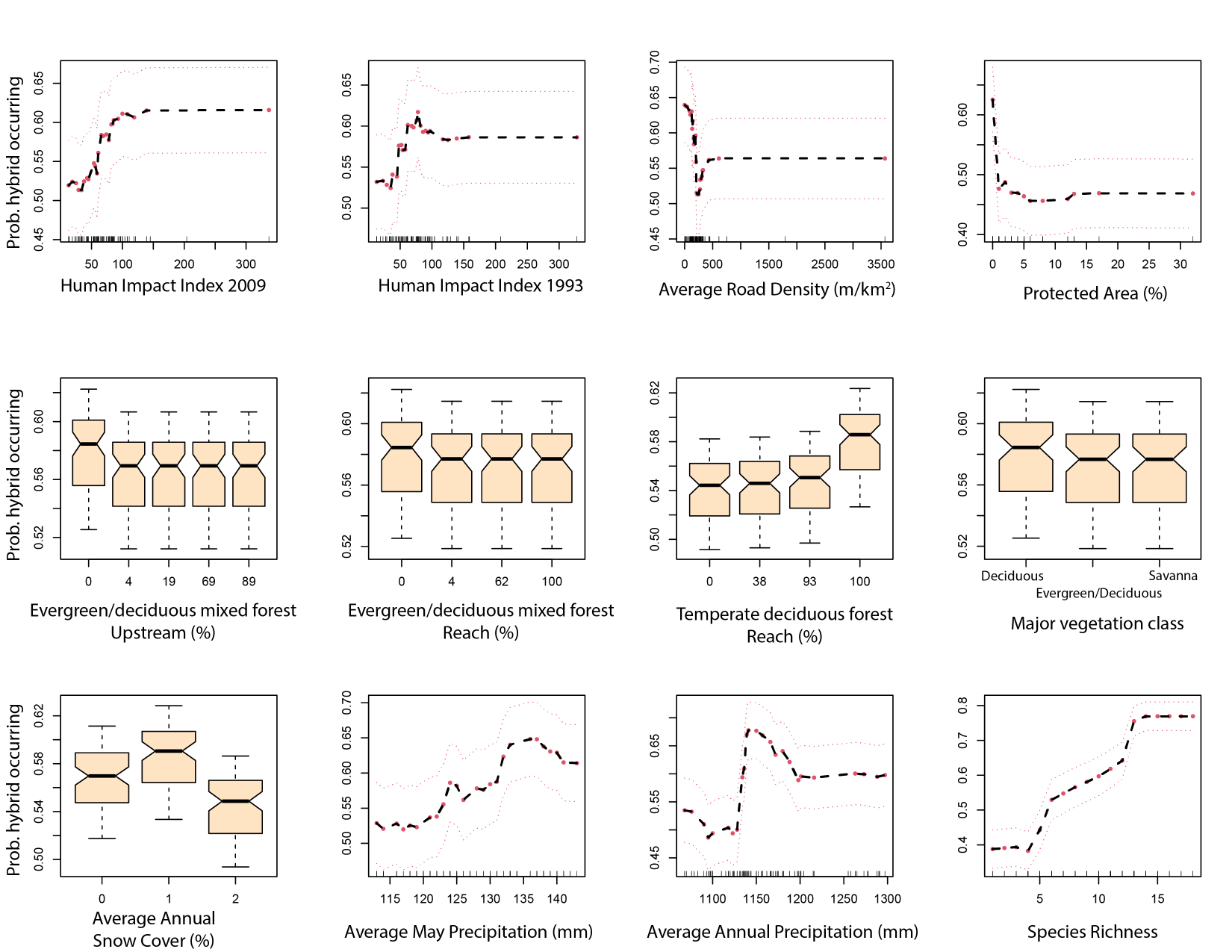

**SUPPLEMENT S25** Six character species codes used in the individual summary table above and throughout code and files.

| **Family** | **Code** | **Common Name** | **Scientific Name** |
| --- | --- | --- | --- |
| Centrarchidae (Sunfish) | LEPMAC | Bluegill Sunfish | *Lepomis macrochirus* |
|  | LEPMEG | Longear Sunfish | *Lepomis megalotis* |
|  | MICDOL | Smallmouth Bass | *Micropterus dolomieu* |
|  | MICSAL | Largemouth Bass | *Micropterus salmoides* |
| Cottidae (Sculpin) | COTCAR | Banded Sculpin | *Cottus carolinae* |
|  | COTHYP | Knobfin Sculpin | *Cottus hypselurus* |
| Fundulidae (Topminnows) | FUNCAT | Northern Studfish | *Fundulus catenatus* |
|  | FUNOLI | Blackspotted Topminnow | *Fundulus olivaceus* |
| Ictaluridae (Catfish) | NOTALB | Ozark Madtom | *Noturus albater* |
|  | NOTEXI | Slender Madtom | *Noturus exilis* |
|  | NOTMAY | Black River Madtom | *Noturus maydeni* |
| Leuciscidae (Minnows) | CAMANO | Central Stoneroller | *Campostoma anomalum* |
|  | CAMOLI | Largescale Stoneroller | *Campostoma oligolepis* |
|  | CHRERY | Southern Redbelly Dace | *Chrosomus erythrogaster* |
|  | CYPGAL | Whitetail Shiner | *Cyprinella galactura* |
|  | CYPWHI | Steelcolor Shiner | *Cyprinella whipplei* |
|  | LUXCHR | Striped Shiner | *Luxilus chrysocephalus* |
|  | LUXPIL | Duskystrip Shiner | *Luxilus pilsbryi* |
|  | LUXZON | Bleeding Shiner | *Luxilus zonatus* |
|  | LYTUMB | Redfin Shiner | *Lythrurus umbratilis* |
|  | NOTBOO | Bigeye Shiner | *Notropis boops* |
|  | NOTNUB | Ozark Minnow | *Notropis nubilus* |
|  | NOTPER | Carmine Shiner | *Notropis percobromus* |
|  | NOTTEL | Telescope Shiner | *Notropis telescopus* |
|  | PIMNOT | Bluntnose Minnow | *Pimephales notatus* |
|  | SEMATR | Creek Chub | *Semotilus atromaculatus* |
| Percidae (Darters) | ETHBLE | Greenside Darter | *Etheostoma blennioides* |
|  | ETHCAE | Rainbow Darter | *Etheostoma caeruleum* |
|  | ETHFLA | Fantail Darter | *Etheostoma flabellare* |
|  | ETHJUL | Yoke Darter | *Etheostoma juliae* |
|  | ETHSPE | Orangethroat Darter | *Etheostoma spectabile* |
|  | ETHUNI | Current Darter | *Etheostoma uniporum* |
|  | ETHZON | Banded Darter | *Etheostoma zonale* |
